# Identification of side effects of COVID-19 drug candidates on embryogenesis using an integrated zebrafish screening platform

**DOI:** 10.1101/2023.06.29.547094

**Authors:** Alexander Ernst, Indre Piragyte, MP Ayisha Marwa, Ngoc Dung Le, Denis Grandgirard, Stephen L. Leib, Andrew Oates, Nadia Mercader

## Abstract

Drug repurposing is an important strategy in COVID-19 treatment, but many clinically approved compounds have not been extensively studied in the context of embryogenesis, thus limiting their administration during pregnancy. Here we used the zebrafish embryo model organism to test the effects of 162 marketed drugs on cardiovascular development. Among the compounds used in the clinic for COVD-19 treatment, we found that Remdesivir led to reduced body size and heart functionality at clinically relevant doses. Ritonavir and Baricitinib showed reduced heart functionality and Molnupiravir and Baricitinib showed effects on embryo activity. Sabizabulin was highly toxic at concentrations only 5 times higher than Cmax and led to a mean mortality of 20% at Cmax. Furthermore, we tested if zebrafish could be used as a model to study inflammatory response in response to spike protein treatment and found that Remdesivir, Ritonavir, Molnupiravir, Baricitinib as well as Sabizabulin counteracted the inflammatory response related gene expression upon SARS-CoV-2 spike protein treatment. Our results show that the zebrafish allows to study immune-modulating properties of COVID-19 compounds and highlights the need to rule out secondary defects of compound treatment on embryogenesis. All results are available on a user friendly web-interface https://share.streamlit.io/alernst/covasc_dataapp/main/CoVasc_DataApp.py that provides a comprehensive overview of all observed phenotypic effects and allows personalized search on specific compounds or group of compounds. Furthermore, the presented platform can be expanded for rapid detection of developmental side effects of new compounds for treatment of COVID-19 and further viral infectious diseases.

**Summary statement:** A zebrafish screening platform assesses side effects on cardiovascular development and behavior of FDA approved drugs used in clinical practice to treat COVID-19 and their immune modulatory effect upon spike protein treatment.

## Introduction

The COVID-19 pandemic demanded the health care system as well as the scientific community to react fast to reduce the spreading and mortality of the disease. Research investigating therapeutics and vaccines has been set in motion with unprecedented speed and the US Food and Drug Administration (FDA) launched several Emergency Use Authorization announcements. The Expanded Access (EA) Program gained importance to promote emergency use of unapproved agents to treat COVID-19 outside of clinical trials. In this regard, drug repurposing became an important research pillar. Indeed, a more general therapeutic treatment beyond vaccination will remain of great clinical importance given the observations that vaccinations provide immunity of a limited duration ^1^ and that newly arising SARS-CoV-2 variants can escape to some extent or completely the immune response obtained through vaccination ^2,3^. One important caveat of drug repurposing is that even though the compounds were previously approved for treatment of other diseases or are under clinical testing, the impact of many of these agents on embryonic development is yet unclear, limiting the access of treatment to pregnant COVID-19 patients.

The zebrafish plays an important role in phenotype-based screening in the field of embryonic and cardiovascular development, drug discovery, and toxicity screenings ^4-8^. In comparison to *in vitro*, cell-based systems, the zebrafish allows to test drugs in the context of an intact vertebrate organism ^9^. With its small size, transparency, rapid external development and high degree of genetic similarity to humans it allows fast assessment of how drugs effect defined developmental stages at tissue and cellular resolution.

The zebrafish heart starts to beat at 25 hours post-fertilization (hpf). The vasculature is initially formed by the dorsal aorta and the cardinal vein from which intersegmental vessels (ISV) sprout at the somite boundaries. The sprouting of these endothelial cells can be used as a proxy for proper angiogenesis and this process is controlled by signaling cascades that are conserved in humans ^10^. Fluorescent myocardial and endothelial reporter lines make imaging straight-forward and scalable when combined with medium-to high-throughput microscopy in 96 well-plates on multiple resolution levels using template-matching algorithms ^11,12^. For example, such a setup has been successfully used for high-throughput identification of small molecules that affect human embryonic vascular development with direct applicability to cancer treatment ^13,14^. Furthermore, the zebrafish was used to identify compounds with cardio-protective function in humans ^15^ and small molecules that promote cardiomyocyte proliferation also found to be effective on mouse cardiomyocytes ^16^. Furthermore, studying the swimming behavior of zebrafish can provide insights into sensory and motor function of developing embryo, and this approach can also be adapted for drug screening in a high-throughput setup ^17,18^.

Here we used the zebrafish model organism to screen for possible side effects on cardiovascular and embryonic development of 162 marketed drugs that are either already used in clinics or being considered as potential treatments for COVID-19. While the effect of some compounds has been previously studied individually in zebrafish, a systematic and comprehensive analysis allowing for comparison between compounds was lacking. We also included the compound Sabizabulin not yet tested in the zebrafish model and currently used as FDA-approved drugs for COVID-19 treatment. We implemented a screening pipeline that includes U-net architectures for automatic image segmentation and analysis, which can be easily scaled up for additional compounds as they become available. The results were uploaded on a straightforward web-based interface that is easily accessible and allow further mining of results. Additionally, we show that the compounds can be efficiently tested for their ability to modulate immune signaling in upon SARS-CoV-2 spike treatment. Our results not only have important implications for the treatment of COVID-19, but also provide valuable insight into the unintended effects of FDA-approved drugs on early development and cardiovascular system formation. Moreover, our pipeline can be adapted for determining the possible effects of further compounds on embryonic development or the cardiovascular system.

## Results

### Establishing a quantitative screening platform to identify drug effects on embryonic and cardiovascular development

One of the earliest initiatives to promote research on drug repurposing to fight COVID-19 was supported by the *Medicine For Malaria Venture (MMV).* The consortium assembled and freely distributed the *Covid Box* for research, comprising a drug library of 160 compounds with known or predicted effects on one or multiple steps of SARS-CoV-2 infection or COVID-19 disease progression. The majority of compounds in the *Covid Box* are categorized as anti-infective agents, that would target biological processes of pathogens (56 agents; 35 % of the library), all the other types would target biological processes in the host (Figure 1a). As an extension of our screening efforts, we also purchased Molnupiravir and Sabizabulin (from MedChemExpress), recently approved agents for the treatment of COVID-19 that successfully inhibit viral replication and transmission from cell to cell, respectively ^19,20^.

**Fig. 1.**
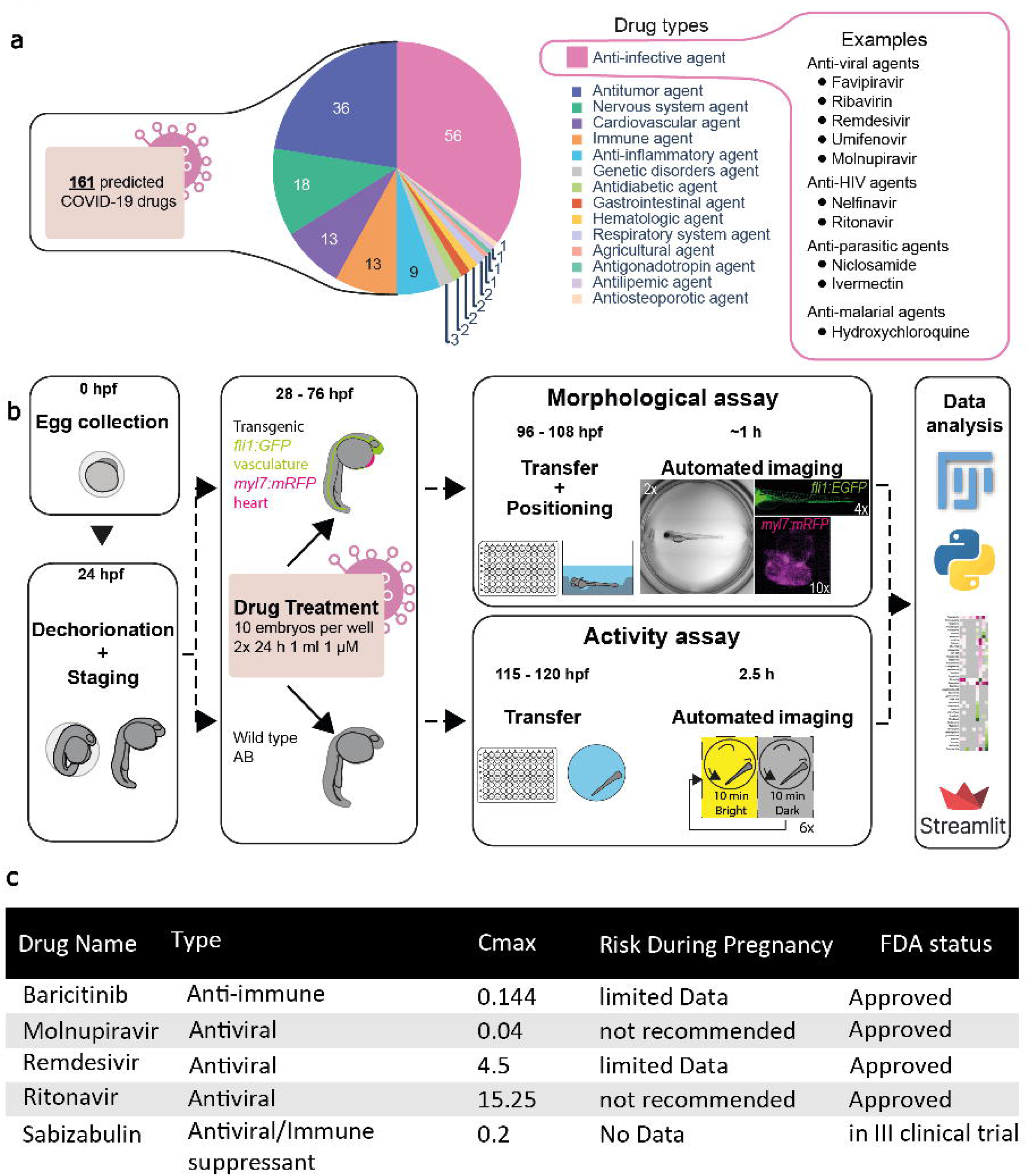
Workflow of zebrafish cardiovascular and behavioral screening setup. **a** Information on type of candidate drugs used. All primary drug indications are listed and a set of anti-infective agents is shown. **b** Schematic representation of the experimental workflow. The time of each step is indicated as hour post fertilization (hpf). For the morphological assay, Tg(*fli1:EGFP*);(*myl7:mRFP*) embryos are used to have fluorescence labeling of the vasculature (green: cytoplasmic GFP in endothelium) and the heart (magenta: membrane RFP in cardiomyocytes). Between 96 and 104 hpf, the anesthetized larvae were transferred to 96 well plates with pre-formed agarose beds by a 3D printed mold. The morphological effects are documented using an automated microscopy platform at three magnification levels with the 2x, 4x and 10x objectives. Shown are representative images at each magnification with brightfield overview (entire well, 2x), trunk vasculature (GFP, 4x) and heart (RFP, 10x). For the behavioral assay, wild type larvae were treated with the drugs and transferred to 96 well plates without anesthesia to swim free in the wells. The swimming of larvae with a defined light-dark exposure was recorded and the tracking data exported. Python and the Fiji software were used to analyze the collected data. The results were presented as heatmaps and in a customized online *Streamlit* data app. **c** A representative table of drugs that are FDA-approved or in III Clinical trial with the indicated C_max_ blood concentrations and risk during pregnancy.

We treated zebrafish embryos at 28 hours post fertilization (hpf) with the compounds dissolved in DMSO (Figure 1b). The baseline concentration for the screening of the drugs was 1 µM, as recommended by the MMV, which was approximately 10x lower than used commonly in zebrafish screens, e.g. ^21-23^. For those compounds also used in the clinical context for COVID-19 treatment we used compounds at C_max_ (Figure 1c). Treatment with each of 162 compounds on 10 zebrafish embryos was replicated two times or more. For Ivermectin and Sabizabulin, as 1 µM concentration elicited severe toxic effects, we lowered the dose. Treated embryos were then subjected to two parallel imaging platforms, that allowed us to observe embryonic and cardiovascular development on the one hand, and behavior on the other. We analyzed the data in a combined workflow in semi-automatic as well as automatic manner to obtain quantitative data for each drug (Figure 1b).

To observe morphological and cardiovascular features, we used the homozygous double transgenic line Tg(*fli1:EGFP*)^y1^;(*myl7:mRFP*)^ko08^ in which the *fli1a* promotor controls expression of the *enhanced green fluorescent protein* encoding gene (*EGFP*) and the *myl7* promoter drives expression of the *membrane-tagged red fluorescent protein* encoding gene (*mRFP*) ^24,25^. At 4 days post fertilization (dpf), larvae were transferred to 96-well plates, prepared with beds of low-melting agarose using a 3D-printed mold and imaged using a High-content Smart Imaging Fluorescent Microscope (Supplementary Fig. S1).

On the other hand, we evaluated swimming behavior, an established method for testing neuroactive compounds (Basnet et al., 2019), using DanioVision^TM^ recording chamber. Tracking the behavior of zebrafish larvae is based on switching between bright and dark phases and can reveal anxiety, vision impairment, muscular weakness and reactivity ^18^ (Supplementary Fig. S2).

### A quarter of tested compounds altered cardiovascular development, embryonic growth, or behavior in the zebrafish model

First, we evaluated the survival and pericardial effusion of the embryos during treatment for both assays (Figure 2a and Supplementary Fig. S3). The mortality rate (MR) was calculated as the average percentage of the larvae (n = 20-60 larvae per drug) that died before imaging in the respective assay. We observed that 7 out of 162 drugs (4.3%) caused a MR over 10% at 1 µM, including Apilimod, Astemizole, Ivermectin, Manidipine, Midostaurin, Niclosamide and Pimozide. Moreover, 18 drugs (about 12.5%) caused pericardial effusion in a quarter or more of the treated larvae. Among them, Astemizole (94.4%), Pimozide (100%) and Ponatinib (100%) were the compounds most frequently leading to pericardial effusion.

**Fig. 2.**
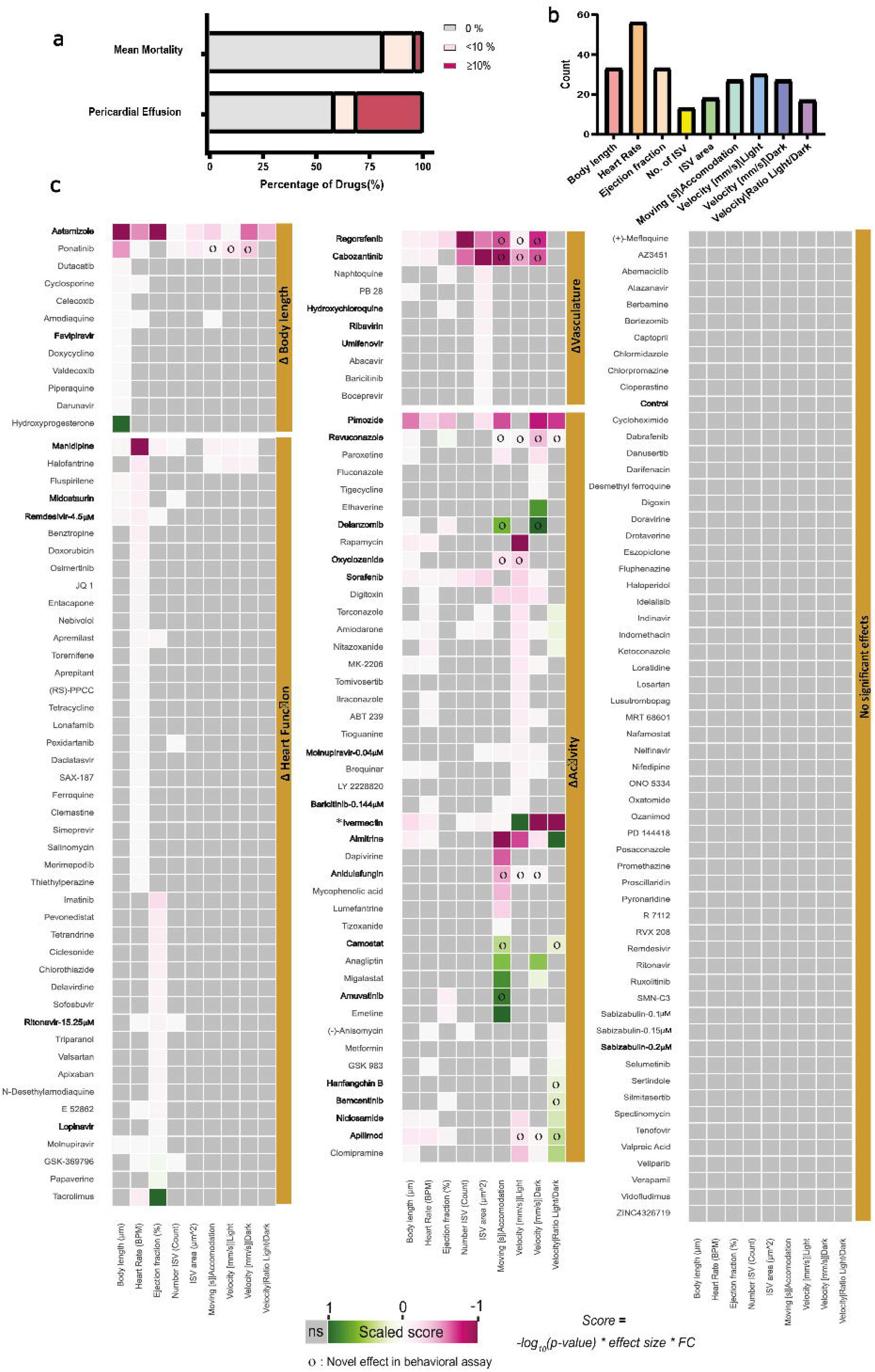
Summary of results obtained for the compounds screening. **a** Bar plot representing the percentage of mortality and pericardial effusion of embryos tested in morphological assay. **b** Bar plot representing percentage of compounds leading to statistically significant alterations in the analyzed parameters. **c** Heat map of the results obtained for the tested compounds. Compound names are listed on the right of the heat map and are allocated to groups: ΔHeart, ΔBody length, ΔVasculature, ΔActivity or “No significant effect”, depending on the results. E.g. a compound that led to high reduction of ejection fraction compared to controls would appear in the group ΔHeart. The formula used to calculate the scores in the heatmap is given at the bottom. The p-value is derived from the Mann-Whitney U test treatment vs. control, the lower the p-value the higher the –log10(p-value). The hedge’s effect size gives an estimation of how strong an effect is. The fold change (FC) shows whether the values are higher or lower than the control and how many times compared to the control. The scores are shown in a relative color scale: green, most positive score; magenta, most negative score; grey, no significance using a Mann-Whitney U test. Shown are the mean values from at least two technical replicates, each replicate is composed of at least 10 embryos per condition. o symbol indicates novel statistically significant observations in behavioral assay. All the compounds are used at 1 μM except when it is mentioned otherwise. Asterisk in Ivermectin refers to the fact that it was used at 1 μM for the morphological assay and at 0.5 μM for the behavioral assay. Compounds mentioned throughout the manuscript are highlighted in bold.

We measured the body length and several parameters to assess cardiovascular development and behavior of larvae and clustered them according to their effect on body length, heart function (rate and ejection fraction), vasculature formation (number and area of ISV) and effect on activity (moving during accommodation, velocity under bright light and dark conditions as well as the ratio of velocity between dark and bright phases) (Figure 2b,c). A final cluster was formed by those compounds that at the tested concentrations did not produce any deviation in the analyzed parameters when compared to controls. We found that 20.3% (33/162) of the drugs significantly altered the size of the larvae. 56 drugs (34.5%) altered the heart rate and 33 drugs (20%) affected the ejection fraction. For assessment of behavior in the presence of the compounds, we scored three phases of the bright/dark locomotion test - accommodation, bright and dark phases - and calculated the ratio between swimming velocity in the dark and bright. Of the drugs tested, 16.6% (27/162) showed alterations in activity during accommodation, with 6 increasing the activity. After a dark accommodation phase, switching the light on causes a startle response in control embryos ^17^. In 18.5% (30/162) of the drug treatments, activity in the bright phase was altered, with 29 compounds causing a reduction in the startling response velocity. In the dark phase, larvae returned to an active state and increased their swimming activity. We found a significant modification of activity in 27 drug treatments, with 23 drugs reducing velocity in the dark compared to controls. To understand whether the difference between activity in the bright and dark phases was proportional to the control, we calculated the logarithmic bright/dark ratio. A reduction in the dark was observed in 66.6% (18/27) of drugs accompanied by a decreased activity in the bright phase. The bright/dark ratio was altered in 10.5% (17/162) of the drug treatments, with 11/17 treatments caused by a reduction of swimming in the bright phase and no or relatively lower reduction in the dark. This could indicate impaired perception of the light stimulus. Of all compounds tested, Ivermectin treatment also caused the most severe effects on swimming behavior, consistent with previous findings ^26^ and Almitrine led to the highest increase in relative swimming velocity (light vs Dark)The effect on swimming behavior of Ponatinib, Regorafenib, Cabozantinib, Ravuconazole, Delanzomib, Oxyclozanide, Anidulafungin, Camostat, Amuvatinib, HanfangchinB, Bemcentinib and Apilimod and Molnupiravir had to our knowledge not been reported before.

We looked closer to those compounds leading to more pronounced defects on development (Figure 3A,B). Astemizole, Pimozide, Ponatinib, and Ivermectin led to the highest reduction in body length. While these compounds had been studied in the zebrafish ^27-30^, size reduction had not been documented. Furthermore, 17α-Hydroxyprogesterone caused a previously not described larval overgrowth. Some of the observed effects on heart rate and ejection fraction had previously been reported, as was the case for Manidipine and Astemizole ^31,32^. However, neither the increase in ejection fraction by Tacrolimus nor the effects of Ravuconazole (an antifungal agent), GSK-369796 (an antimalarial compound) Amuvatinib and Regorafenib have to our knowledge not been reported before. The Tyrosine kinase inhibitors Regorafenib, Cabozantinib, and Sorafenib revealed a strong inhibition of ISV formation, as reported before, and thus served as a positive control for the experimental set up ^33-35^. However, the negative effects on ISV area by the compounds Astemizole and Pimozide had not been reported before.

**Fig. 3.**
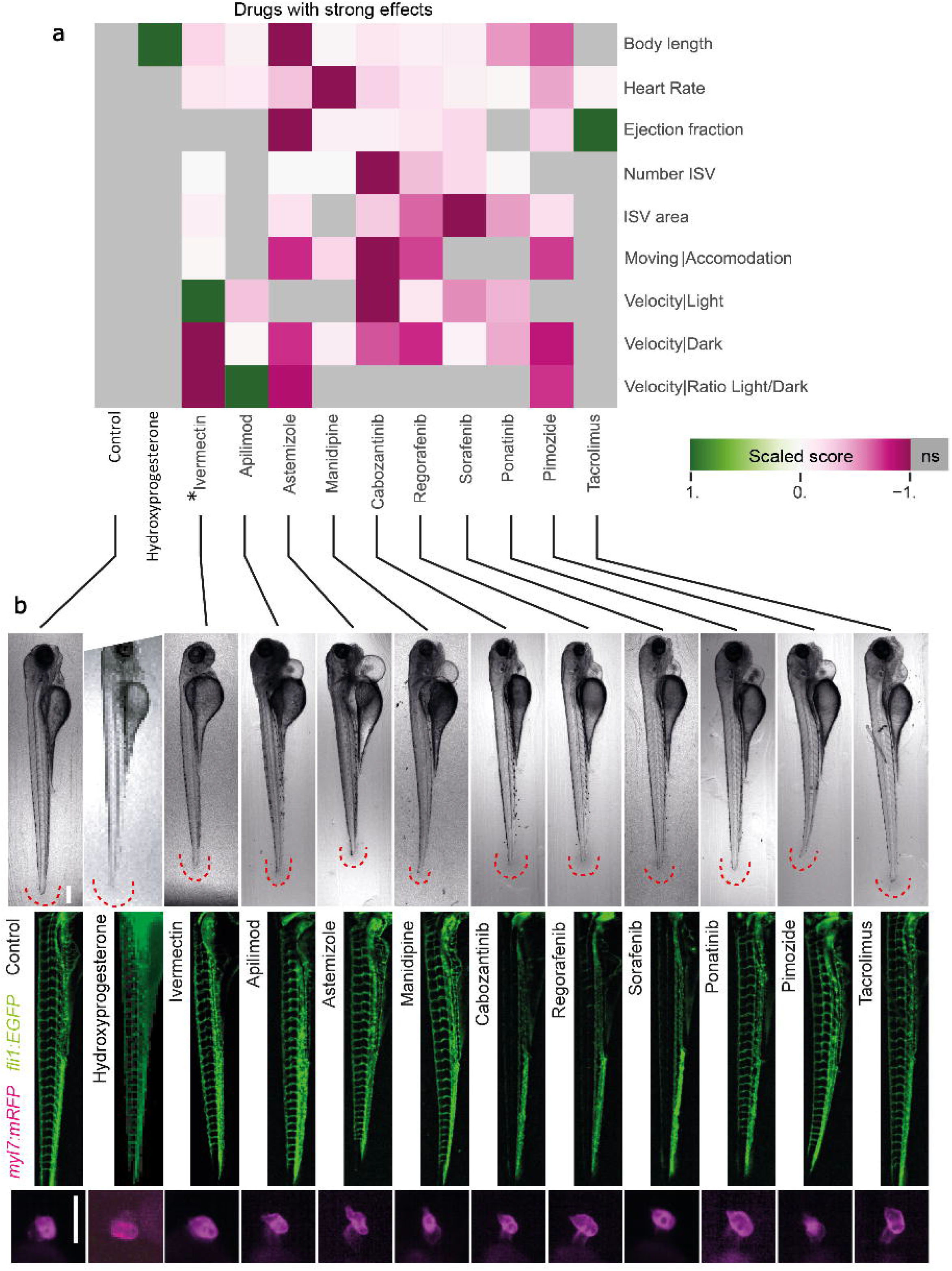
Selection of compounds eliciting severe effects on zebrafish embryonic development. **a** Results of 11 drug treatments leading to strong phenotypic alterations. Heatmap showing the highest positive score in dark green and the lowest negative score in dark magenta. Each measurement is individually scaled. Grey color indicates no significance according to the Mann-Whitney U test. Shown are the mean values from at least two technical replicates, each replicate is composed of at least 10 embryos per condition. Boxplots with individual data points for the morphology assay can be found in Fig. S4 and the behavioral assay Fig. S5. **b** Example images of the control and treatments with visible effects are shown in brightfield, vasculature Tg(*fli1:EGFP*) and heart Tg(*myl7:mRFP*). Brightfield and GFP images are shown as sobel-projections. The dynamic range of GFP images was homogenized using the DEVILS Fiji-plugin. Scale bars: 250 μm.

In conclusion, the combination of results from two established assays for 162 clinically relevant compounds indicated that roughly one fourth of these drugs altered cardiovascular development, overall embryonic growth, or behavior in the zebrafish model.

### Effect of compounds studied in the context of COVID-19

We focused at the impact of those compounds with the highest interest for COVID-19 treatment on cardiovascular development and larval activity in zebrafish. Ivermectin and Hydroxychloroquine were compounds repurposed for COVID-19 treatment in the initial stage of the pandemic but their use was discontinued ^36-38^. Ivermectin was highly toxic at the concentration of 1 µM. The Effect Score (ES) for body length, heart rate, as well as ISV number and area were significantly reduced upon 1 µM Ivermectin treatment (Figures 2c, 3 and Supplementary Figs. S3-6). At 4 dpf the heart rate was drastically reduced and all embryos died between 4.5 and 5 dpf. At a lower dose of Ivermectin (0.5 µM) mortality disappeared, but we still observed effects on larval behavior (Figures 2b and 3). The results are in line with reported toxicity of Ivermectin in zebrafish and its side effects in human patients^30,36^.

Hydroxychloroquine treatment showed significant reductions of ejection fraction and ISV area (Figures 2c, 3 and Supplementary Fig. S4). The effects of Hydroxychloroquine are in line with mild cardiac phenotypes, observed in neonates after treatment during pregnancy and increased cardiovascular mortality ^39^.

Clinical trials on Favipavir, Ribavirin, Umifenovir and Lopinavir have been started ^40^ but are not FDA approved for COVID-19 treatment. Favipiravir had mild effect on body length and Lopinavir reduced ejection fraction (Figure 2b). We found that Ribavirin and Umifenovir treatment mildly reduced the ISV area (Figure 2b). While the effects observed for these compounds showed statistical significance, the amplitude of these effects was very low.

Remdesivir, Molnupiravir ^19^, Paxlovid (Nirmatrelvir and Ritonavir) ^41^, and Baricitinib are all used in the clinics and were approved by the FDA for COVID-19 treatment ^42,43^. Sabizabulin has also been recommended for COVID-19 treatment ^20^. We analyzed effect of compounds Remdesivir, Ritonavir, Baricitinib, Molnupiravir, and Sabizabulin on our zebrafish assay at the highest concentration detected in blood plasma in humans (C_max_)^44-47^ (Figure 4b and Supplementary Fig. S4c). Sabizabulin treatment still resulted in a 20% mortality at its C _max_ (0.2μM) (Figure 4a). Those larvae that survived, did however not present any cardiovascular or behavioral defects (Figure 4b). At C_max_ (4.5 μM) Remdesivir led to decrease in heart rate, ejection fraction and in body size of the embryo (Figure 4b). For Ritonavir, C_max_ was considerably higher than the tested concentrations (15.25 μM). We found that this altered the ejection fraction and heart rate (Figure 4b). Baricitinib at C _max_ (0.144 μM) led to 18.3 % mortality and in those embryos that survived caused alterations in heart rate and motility (Figures 4a and b). Despite a C_max_ of only 0.04 μM, at this concentration Molnupiravir still altered swimming behavior and a decrease in body size.

**Fig. 4.**
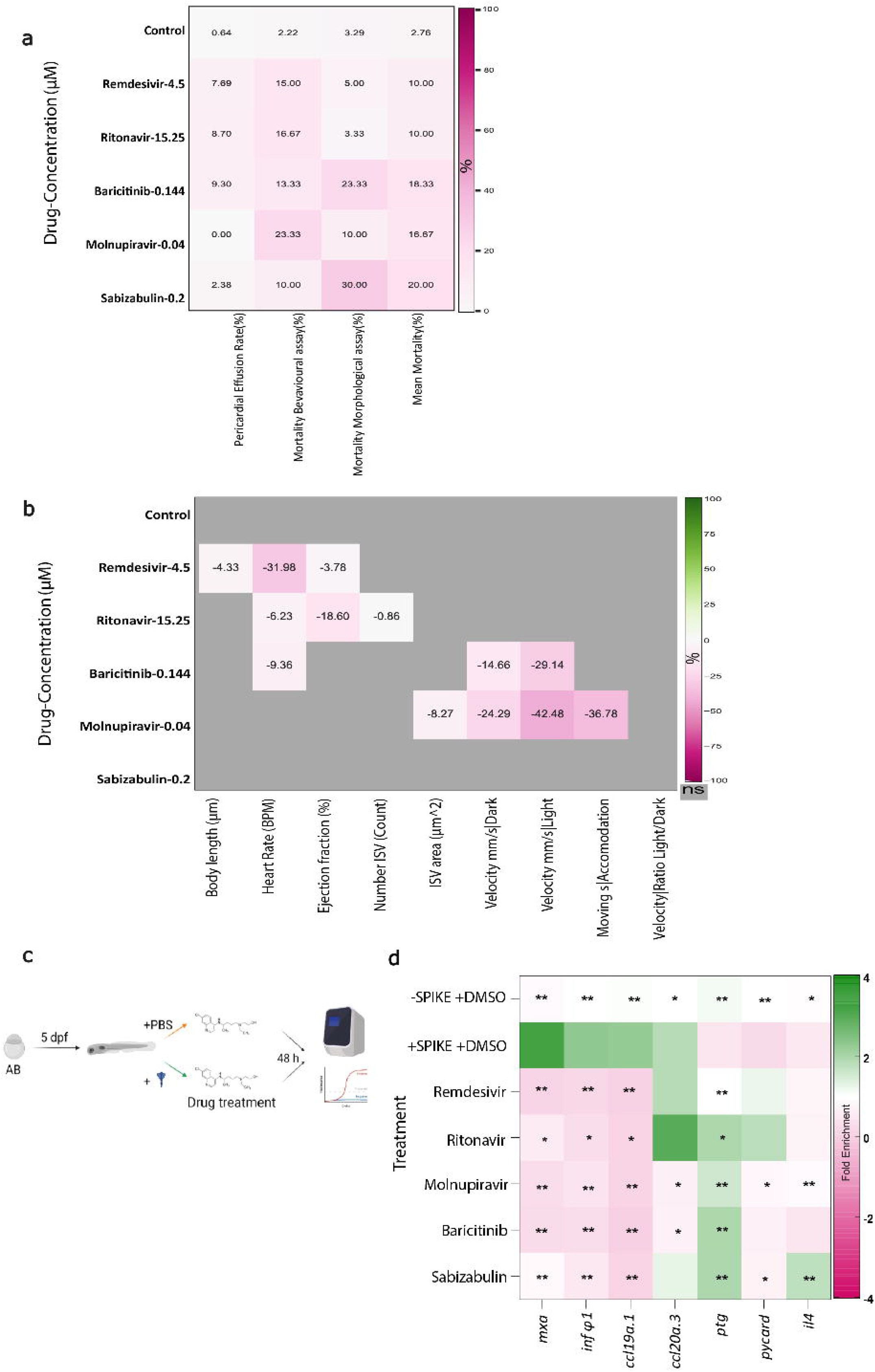
Selected compounds at clinically relevant concentrations result in mild modulation of embryonic development and immune suppression in response to SARS-CoV-2 spike treatment. **a** A heatmap showing Percentage of Pericardial effusion and Mortality of the selected drugs at their clinically relevant concentrations. Mortality rates for Morphological Assay(with PTU) and Behavioral Assay (without PTU) are plotted separately followed by a column of mean mortality rate. Dark magenta shows the highest effect. **b** The relative percentage effect of these selected drugs at their clinically relevant concentrations are shown in the heatmap. The highest positive effect is shown in dark green and the lowest negative effect is dark magenta. Grey color indicates no significance according to the Mann-Whitney U test. Shown are the percentage from median values from at three technical replicates for drugs except for Remdesivir, that was tested in two replicates, each replicate is composed of at least 10 embryos per condition during drug treatment. **c** Scheme of larvae treatment with SARS-CoV-2 spike protein and selected drugs. **d** qPCR results represented as a heatmap representing fold change with respect to treatment with DMSO without spike (-SPIKE +DMSO). The upregulation is shown in dark green and the downregulation in dark magenta. Significance calculated between DMSO with SPIKE-protein treated group (+SPIKE+DMSO) and the rest. Each color box represents a mean of 6 biological replicates of 5 embryo pools (n=6, Mann-Whitney U-test treatment vs. + spike + DMSO; * p< 0.05, ** p<0.01, *** p< 0.001).

In sum, at clinically relevant concentrations, all five compounds used in the clinical context led to alterations in embryonic development.

### Drug candidates modulate zebrafish immune response to SARS-CoV-2 spike protein treatment

Zebrafish possess many similarities in the innate immune response to that of mammals ^48^. While successful virus amplification was not observed in wild type strains, SARS-CoV-2 spike treatment causes temporal immune response in zebrafish embryos and adult fish ^49-51^. We decided to explore the effect of drugs selected for C_max_ analysis on the modulation of the inflammatory and immune response gene expression profile (Figure 1c). We subjected 5 dpf larvae to recombinant SARS-CoV-2 spike protein and treated them with Remdesivir, Ritonavir, Molnupiravir, Baricitinib, Sabizabulin or DMSO for 48 h (Figure 4c).

First, RT-qPCR confirmed that after spike treatment there is a modulation of several immune response genes compared with a control group of larvae not subjected to spike protein administration (Supplementary Fig. S6). We observed a more than three-fold upregulation of the antiviral immune response genes *mxa*. Other upregulated pro-inflammatory immune response genes were type I interferon *infΦ1,* chemokines *ccl19a.1* and *ccl20a.3*. On the other hand, *ptg* and *pycard* but not *caspa*, all involved in inflammasome pathway ^50^, along with *il4/13b*, an anti-inflammatory gene ^52^, were downregulated.

Next, we performed co-treatment of larvae with spike protein and the FDA-approved drugs at C_max_. Then, we tested the effect of the compounds on alteration of gene expression of the panel of genes, in which we had observed a significant response upon spike protein treatment (Figure 4d). All compounds successfully reduced *mxa*, *infΦ1*, and *ccl19* expression to the levels equivalent or even lower than untreated controls. Baricitinib and Molnupiravir treatments also led to significantly reduced *ccl20a.3* levels. The drug treatments also affected inflammasome pathway genes; *ptg* was upregulated to control levels by Remdesivir and Molnupiravir, while other drugs led to four-fold upregulation. Molnupiravir and Sabizabulin also successfully downregulated *il4* expression. Of note, we also observed that in the absence of spike protein, drug treatments were already able to alter the expression of some of the immune response genes (Supplementary Fig. S7). Of all five drugs, Remdesivir had the broadest effect on inflammatory genes in the absence of spike protein treatment. Overall, these results indicate that the zebrafish model is useful not only for screening phenotypic and behavioral alterations mediated by drug treatment, but also to assess compound effectiveness in attenuating the immune response after SARS-CoV-2 spike protein exposure. It also highlights that at C_max_ clinically relevant compounds can lead to alterations of embryonic development in the zebrafish.

### Data access via an online data app

In order to facilitate data mining of this large dataset we created an online data app using the open-source Streamlit app framework (Fig. 5 and accessible via this link: https://share.streamlit.io/alernst/covasc_dataapp/main/CoVasc_DataApp.py. The online data application allows to access raw measurements as well as batch corrected data and provides access example images for each treatment according to one’s individual interests.

**Fig. 5.**
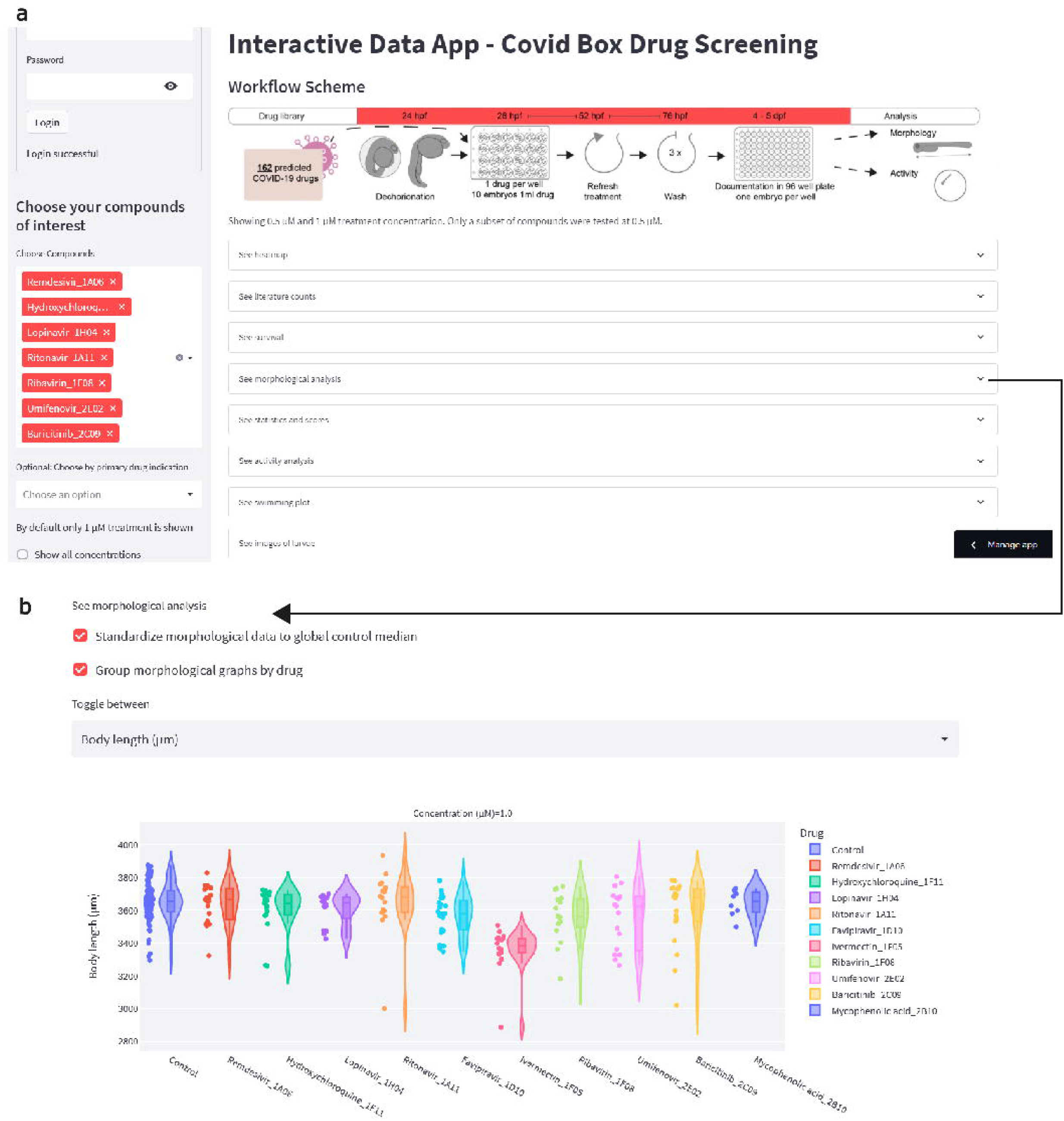
Generation of a Web-based Data App. **a** The data app overview window shows which information can be found in the app. A heatmap with the previously described scores, results of the literature search, survival data, morphological analysis, activity analysis and images of the larvae. **b** Example tab opened, showing interactive violin plots for the larval length, which allows to see the distribution and individual data points. The checkboxes allow to view the corrected or raw data batch, and replicates can be grouped or shown individually. https://share.streamlit.io/alernst/covasc_dataapp/main/CoVasc_DataApp.py

Here an overview of the functionality (Figure 5a): on the left side, we allocated a side bar to select a specific tested compound or group of compounds. In the main window, tabs were created to allow visualization of an overview heat map, literature data, mortality of the treatments, morphological analysis and behavioral analysis. Additionally, we chose representative example images for each drug treatment group in each experimental replicate. An overview in brightfield and GFP fluorescence was provided in the last tab. Within the tabs we gave options to visualize the data (Figure 5b). We provided options to show batch corrected data (check box: “Standardize to global median”) and to visualize individual experiments with the respective control or to group replicates by the treatment.

By providing the data app, we enable better insight into the data and facilitate visualization of all screening results beyond the ones shown in the figures. Furthermore, this app could be considered as a layout to be implemented for other screening projects.

## Discussion

Our motivation to perform this study was to support drug repurposing efforts in the context of the COVID-19 therapy, however, given that FDA approved compounds have been tested, the assay provides information that can be useful beyond this disease. Despite of the rapid progress of drug development and many clinical trials, the quest on the search of an ideal drug therapy against COVID-19 continues. Even though several compounds have been approved for COVID-19 treatment, most of these drugs have not undergone thorough investigation for pregnant individuals ^53-56^, or are not indicated during pregnancy without providing further information, as is the case for Remdesivir, Molnupiravir, and Ritonavir ^57^. Here we report the use of the zebrafish model to assess effectiveness and side effects on cardiovascular development of compounds repurposed for COVID-19 treatment. From the compounds that are currently used in the clinic Baricitinib, Remdesivir, Ritonavir had been studied for the induction of embryonic developmental defects in the zebrafish, but not in the context of cardiovascular development, and not all at concentrations corresponding to C_max_ ^58,59^. To our knowledge the effect of Molnupiravir and Sabizabulin on zebrafish development have not been studied to date.

We confirmed that Ivermectin severely affected embryonic development. Indeed, prevention and treatment of COVID-19 by Ivermectin has been warned against by the FDA due to lack of evidence for therapeutic benefits and adverse effects already during the pandemic (FDA, 2021). The compound Hydroxychloroquine caused mild defects on the cardiovascular system. While Hydroxychloroquine’s use in COVID19 treatment has been discontinued, it is still used for treatment of Malaria. Thus, further work elucidating possible side effects on embryonic development are still of relevance. Regarding compounds currently used for COVID-19 treatment, we observed about 20% mortality at C_max_ for Sabizabulin. This microtubule assembly disruptor has entered in a III Clinical trial stage and was shown to reduce deaths of severe hospitalized patients by almost 25%. Our results call for further work to study contraindication of Sabizabulin during pregnancy. Molnupiravir and Paxlovid (Nirmatrelvir and Ritonavir) ^41^ were approved by the FDA, but the approval is still based on little clinical experience in particular regarding patients with health conditions ^42^. Up to this point, we studied only individual compounds and no combination treatments like Paxlovid. Comparing the results of two viral-replication inhibitors Remdesivir and Molnupiravir, both treatments seem to be inappropriate as severe adverse effects were observed. Ritonavir treatments caused overall little embryonic alterations, but led to significant impairment of the heart functionality and Baricitinib treatment asl affected heart rate and embryonic behavior. Overall, all five compounds let to a larger or lesser extent to alterations in embryonic development.

We have also established a workflow to treat 5 dpf embryos with SARS-CoV-2 SPIKE protein to assess whether any of the relevant candidates could affect immune signaling induced by spike treatment. All five drugs tested for modulation of immune response after SARS-CoV-2 SPIKE treatment showed a significant reduction in inflammatory gene expression, namely *mxa*, *infΦ1*, and *ccl19a.1*. Our results show that the zebrafish model allows not only to study the immunogenic nature of Sars-Cov-2 viral particles, that can occur in the absence of infection (Tykalska et al 2022) but also to address how compounds modulate such an inflammatory/immunogenic response. We did observe some differences in the gene expression response to spike protein compared to (Tykalska et al 2022) with no significant effect of spike treatment on *il1b*, *tnfa*, *nfkb1*, *cxcl8a*, *ifnγ*, *caspa* and *il10* expression. This is likely due to differences in the embryonic stage used for the assay and due to the use of whole embryos in our study.

Even though several lines of evidence point to the validity of the zebrafish model to study physiological effect of chemical compounds as a first hint towards its effect in mammals, including humans ^6^, species-specific effects such as differences in target conservation cannot be fully excluded. A further limitation is the route of administration. In comparison to treatments in mammals, zebrafish embryos are commonly treated by immersion. This treatment procedure may limit the estimation of the actual dose received by the organism due to limited penetration. Solubility in water may be another limiting factor. Compared to other zebrafish screenings ^60,61^, the used standard concentration of 1 µM is relatively low and should thus cause more specific effects and less general toxicity. Additionally, less issues regarding solubility can be expected. However, the concentration of individual compounds might vary massively if compared to the concentrations used in clinical practice. Indeed, effects seen upon low drug dosage were also reported to have an effect in mammals, indicating translational potential ^62^. Therefore, to provide more translationally meaningful results we included drug treatment on zebrafish using concentrations relevant to clinical studies (C_max_).

Automated screening setups produce large amounts of image data, which have to be analyzed in a standardized workflow. Machine learning has become an important tool in image analysis and is based on providing examples of a specific target structures to the computer ^63^. In particular, convolutional neural networks nowadays outperform most conventional image analysis tools. In biomedical and medical image segmentation, U-Net architectures prove to be highly versatile ^64^. Here we provide a deep learning model for fast and precise segmentation of embryonic zebrafish hearts and ISVs between 4-5 dpf. The model can be used directly for future screening assays, using the same imaging modalities or can be employed for transfer-learning to reduce the amount of necessary training data.

Besides the code for the U-Net training and analysis, we provide with this work a variety of tools to facilitate future zebrafish developmental screenings. We include multiple Jupyter notebooks and Python functions for image and data analysis, ImageJ-macros for image processing and semi-automatic analysis, the code to perform a systematic literature search, a customized template for a 3D-printed mold to position larvae in 96-well plates and a data app including the full code to allow individual data visualization.

While the herein tested compounds are overall well-studied drugs used in advanced clinical trials and already on the market, our screening assay can also be applied at earlier stages of drug development: before or in parallel to entering clinical trials. The presented screening pipeline requires eight days for a single researcher for 30-40 drugs to be tested at a given concentration. The steps included are: setup of the fish crosses until staging (three days), drug treatment (two days), imaging of morphology (one day) and behavior (one day) and analysis (one day). Future work can aim at reducing the time for analysis by fully implementing deep learning for all analyzed parameters. We avoided such an approach to correlate results from two sources, involving easy and fast counting in a semi-automatic approach and using automatic U-Net based segmentation for the more time-consuming parts. Overall, these two approaches combined with batch correction and scoring delivered highly coherent results pointing out relevant effects on embryonic development.

Many medicines are still insufficiently studied in the context of development, posing serious limitations in the treatment of pregnant women. We envisage that pipelines using the zebrafish model as the one presented in this study will be of interest to not only in the context of COVID-19 treatment but also for characterization of side effects on embryonic cardiovascular development of further treatments.

## Material and methods

### Zebrafish husbandry and studies on embryos

Experiments were performed with zebrafish (*Danio rerio*) embryos and larva at Institute of Anatomy (National License Number 35) and Institut für Infektionskrankeiten (National Licence Number 113) from the University of Bern. Adult fish needed for breeding were maintained at the Institute of Anatomy and raised and maintained at maximal 5 fish/l with the following environmental conditions: 27.5-28°C, with 14 h of light and 10 h of dark, 650-700 μs/cm, pH 7.5 and 10% of water exchange daily. Animal studies were approved by the Animal Care and Experimentation Committee of the Canton of Bern, Switzerland (license BE27/2021). All experiments were performed in accordance with the guidelines and regulations approved by the Animal Care and Experimentation Committee of the Canton of Bern, Switzerland. The study was designed in accordance with ARRIVE guidelines. For larvae > 5 dpf used for qPCR analysis, no animals were excluded. Used larvae were not randomized. Confounders were minimized by using as controls larvae from the same clutch as those used for drug treatments. The experimenter was aware of the group allocation at all stages of the experiment. The number of treated and imaged embryos for each experiment and drug is documented in Supplementary Table 1.

### Breeding and staging

Adult zebrafish (wild type or transgenic) were kept in family breeding tanks overnight, male and female fish separated by a transparent screen. For the morphology assay homozygous Tg(*fli1a:GFP*)*^y1Tg^* and Tg(*myl7:mRFP*)*^ko08Tg^*were crossed to obtain heterozygous double transgenic Tg(*fli1a:GFP*)*^y1Tg^;(myl7:mRFP*)*^ko08Tg^*. For behavioral studies, we used zebrafish from the wildtype AB strain. To initiate synchronous breeding the screens of all breeding tanks were removed at the same time and subsequently eggs were collected within 30 min and kept in E3 medium (5 mM NaCl, 0.17 mM KCl, 0.33 mM CaCl_2_, 0.33 mM MgSO_4_) with Methylene Blue (10^-5^%). Eggs from different breeding tanks were collected and then split into petri dishes at equal density. Confounders were minimized by taking controls from the same clutch as the experimental group. 30 min after egg collection unfertilized eggs and those that did not transition to two-cell stage, were removed. The next day, at 24 h after removing the screens, chorions were removed by incubating 2 mg/ml Pronase in E3 medium for about 3 min until gentle shaking of the petridish freed the larvae from the chorion.

### Pharmacological treatment

Larvae were staged according to ^65^ before starting the drug treatments. Then, 28 hpf embryos were transferred to 24 well plates, 10 embryos per well in 2 ml E3 medium. A new 24 well plate was prepared with 975 µl treatment solution (E3, 5 mM HEPES-Buffer with or without 0.003% 1-phenyl-2-thiourea, PTU, 0.25% DMSO) containing the dissolved drug. Embryos were transferred in 25 µL of medium, to reach 1 µM drug concentration. For one experiment, typically 10 drugs were tested and two wells were kept with control embryos (treatment solution as described above with only DMSO). A full list of experimental replicates can be found in Supplemental Table 1.

The well plate was covered from light and placed into the incubator at 28°C. After 24 h of incubation the drug solution was replaced by fresh solution. After additional 24 h the embryos were washed three times with 2 ml E3 medium (with or without PTU) and kept in the respective medium until the effects are recorded.

For RNA extraction, groups 5 dpf AB wild type zebrafish were placed in 24 well plates. Each well contained 1 mL of fish water with 0.1-15.2 µM drug (fish water, 5 mM HEPES-Buffer, 0.25% DMSO) (refer to Fig. 1C) in the presence or absence of SARS-CoV-2 spike protein (5ng/mL; GenScript). Controls were kept in the treatment solution with only DMSO. The plates were kept in the incubator at 28°C for 48h without medium change. Before the collection for RNA, zebrafish were euthanized with 0.16% Tricaine and washed with PBS. We have chosen to use 5 larvae per each treatment, as this enables to extract sufficient RNA for cDNA production (200-300 ng of RNA was obtained per larvae). We also used GPower analysis to predict how many biological replicates coming from different parents we would need for each treatment (n=6). In total 360 larvae at 7 dpf were used for this experiment.

### Preparation of 96 well plates with 3D printed mold

For lateral positioning of larval zebrafish in 96 well plates (Greiner) we used either a 3D printed mold provided by Acquifer Germany or a custom-made 3D print (Template: https://github.com/Alernst/CoVasc_DrugScreen/tree/main/3D_print). Using a multipipette, 70 µl of melted 1.5% low melting agarose (Promega) was injected into each well. Eventually occurring bubbles were removed using a metal needle. The template mold was carefully inserted in the plate and the plate with the mold was placed at 4°C for 30 min. Subsequently, the mold was pulled out and the plate with agarose beds was ready to use.

### Organismal and cardiovascular development assay

Up to four experiments were run per day, for this reason larvae were taken between 96 and 104 hpf. Each experiment was compared to its own control group with the same embryonic stage. Embryos that were harmed due to handling were excluded from the assay. Between 7-10 embryos were taken per experimental replicate per drug. Embryos were transferred from the 24-well plate into a 12 well plate for easier handling with 4 ml volume per well. For anesthesia, tricaine was added at 0.16 mg/ml (pH 7) and the larvae were immediately transferred to the prepared 96 well plate with agarose beds using a cut yellow pipette tip with 70 µl volume, resulting in a final concentration of 0.08 mg/ml tricaine.

The imaging of the larvae in the 96-well plate was performed using an automated microscopy platform (Acquifer Imaging Machine, Acquifer Germany). Initially, an overview image of each entire well was taken with the 2x objective (pre-scan). From all the brightfield and green fluorescence channel overview images (11 Z-planes), we chose a reference image of the trunk and the head. This image served as a template. Subsequently, an implemented algorithm (template matching) was used to detect trunk and head region of the rest of larvae. After successful detection, a 4x objective was used to acquire Z-stacks (25 Z-planes) of all trunks in brightfield and green fluorescence channel. Next, the 10x objective was used to acquire a time series of 100 frames of the head region focused on the heart. For initial screening experiments, 300 frames in brightfield and red fluorescence channel were acquired.

### Semi-automated image analysis

The analysis was performed using the Fiji-software. A customized ImageJ-macro (https://github.com/Alernst/CoVasc_DrugScreen/blob/main/macros_imagej/AIM_Semiauto_Analysi s.ijm) was written to sequentially open the images of the dataset without showing the treatment group. The macro guides the user through the workflow. At each step requiring manual interaction a dialog window shows the instructions which action to perform. 1) To count ISVs draw a line crossing all ISVs in the green fluorescence channel. The macro creates a kymograph along the Z-axis and detect the local maxima in the kymograph. The user is asked to supervise the detected maxima. 2) Based on the shape of the pericardial cavity, a decision is taken whether cardiac pericardial effusion is present or not. 3) Draw a straight line along the anterior-posterior axis of the larva from head to tail to measure the linear axis length, following recommendations ^66^. 4) The macro opens the heart time series, performs a maximum intensity projection along the time axis, detects the heart in the T-projection by thresholding. The macro suggests a region of interest (ROI) that contains the heart, if this ROI is misplaced, the user can correct the location. The macro now creates a perpendicular line in the center of the lower edge of the ROI, creates a kymograph and detects the local maxima, which needs supervision by the user. All measurements were summarized in a result table containing embryo length, heart rate, ISV number and presence of pericardial effusion. Only after the semi-automatic steps, the drugs were assigned to the analyzed data to ensure an unbiased analysis.

### Automated image analysis

The second part of the analysis was performed using Python via Jupyter notebooks. The Jupyter notebook contains each step of the analysis and can be found in the repository, including all required packages (https://github.com/Alernst/CoVasc_DrugScreen/tree/main/Deep_Learning). The aim was to perform a segmentation of the ISVs, similar to ^67^, and the heart distinguishing between ventricle and atrium, comparable to ^68^.

The core components for segmentation are two newly trained convolutional neural network with a U-Net architecture. The first model, now called IsvSeg, was trained to perform a binary segmentation to distinguish background from ISVs, 30 maximum intensity projections (2048×2048) were manually anotated and a data augmentation was performed, also out-of-focus data were included in the training. The second model for multiclass annotation, named HeartSeg, was trained to segment the ventricle, atrium and background. For training, 348 images (512×512) were manually labelled and out-of-focus data were again also included in the training. The model was tested on 96 new set of images and was evaluated with an unweighted dice coefficient of 0.902.

The HeartSeg model was applied to 100 frames of each heart time series in the red fluorescence channel. To increase the quality of the data, small objects under a threshold area were removed and if any frame did not contain a segmentation of the ventricle, the entire dataset was removed. To calculate the ejection fraction the ventricle area at each time point of the times series was extracted from the segmentation. A peak detection algorithm was used to identify all systoles and diastoles. Due to the shape of the embryonic fish heart the 2D area can be used to calculate the ejection fraction ^69^. A median systole and diastole area was calculated per embryo, which was used to obtain the ejection fraction for each larva, using the formula below:

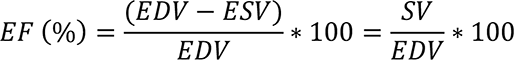

Alterations from the standard experimental procedure in a subset of experiments are explained in the comment section of Supplementary Table 1.

The IsvSeg model was applied to maximum intensity projections of all 4x images of the green fluorescence channel. We measured each ISV’s major (length) and minor axis (width), using the functions included in SciKit image, and calculated a median for each larva. The median ISV area was estimated by multiplying median width and length.

As a final step, we combined the information related to ISV area, ejection fraction, larva length, heart rate, ISV number and pericardial effusion for further meta-analysis (Figure 2c).

### Behavioral analysis

To measure the activity and potential impact of drugs on behavioral patterns of larval zebrafish we used the DanioVision^TM^ recording chamber (Noldus Inc., Wageningen, Netherlands). Between 115– 120 hpf, the previously treated wild-type larvae were placed individually in the wells of a 96 well plate with 200 µl E3 medium pre-warmed to 28°C. No anesthesia or fixation was applied to allow free swimming in the well. The plate was mounted in the DanioVision^TM^ recording chamber. A programmed light cycle is automatically switching on and off the light inside of the recording chamber. Then, larvae had 30 min of accommodation time in the dark, followed by 6 cycles of 10 min dark and 10 min bright periods. During these 2.5 h in total, the larvae were recorded. Larvae tracking upon light stimulation was carried out using the EthoVision XT^TM^ software (Fig. S2). The screening analysis was performed using the 1 min binned images exported from the DanioVision^TM^ recording chamber as xlsx-file. Prior to analysis, incorrectly tracked larvae were excluded by visual inspection of the movies. A table was generated to annotate bright and dark phases as applied in the experiment. All calculations with detailed description were made in Jupyter notebooks which can be accessed on (https://github.com/Alernst/CoVasc_DrugScreen/blob/main/CoVasc_Data_Update.ipynb). From the annotated data a median was calculated for the swimming velocity, moving duration and the distance moved in dark and bright. The logarithmic bright/dark ratio was calculated by dividing velocity in the bright by velocity in the dark and calculating the natural logarithm. To handle zero values, 0.001 was added to both values.

### qPCR

RNA extraction of 7 dpf larvae (5 dpf +2 days of drug treatment) in batches of 5 individuals was performed using TriZol reagent (Sigma-Aldrich) and RNA purification kit (Zymo). cDNA was generated using Maxima First Strand cDNA synthesis kit (Thermo Fisher Scientific). RT-qPCR analysis was performed using PowerUp SYBR Green Master Mix (Thermo Fisher Scientific) and following primers:

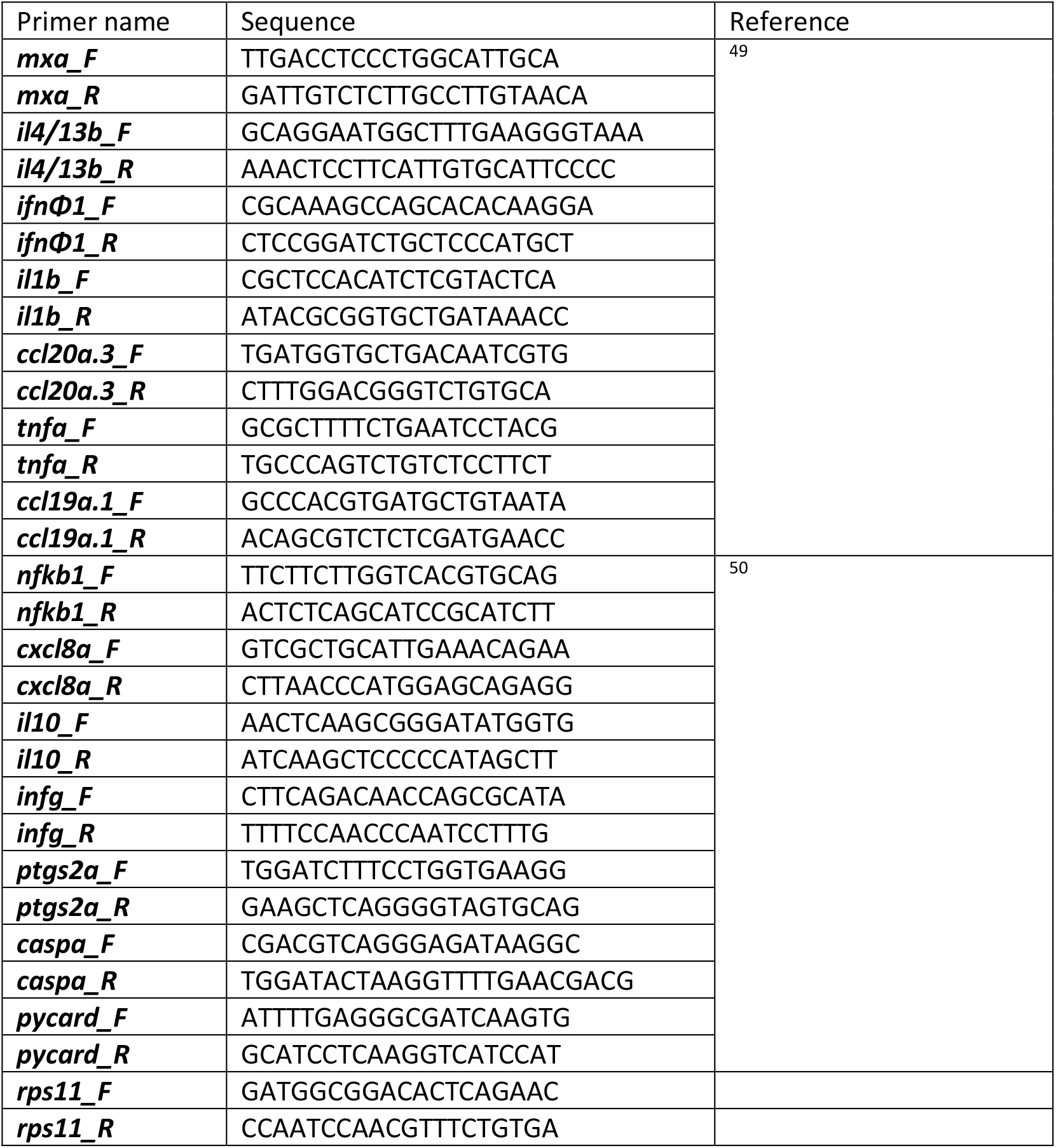

qPCR was performed on an Applied Biosystems QuantStudio 6 Pro System. 6 replicates, each comprising a pool of 5 embryos were performed.

### Statistics and batch correction

All results from Fiji and Python were saved as csv-files and subsequently merged to a single data table. The Mann-Whitney U test was applied to compare each treatment group to the control group of the same experiment (acquired on the same 96 well plate). Additionally, we calculated the effect size and the relative difference of the medians compared to the control. To evaluate multiple experiments and replicates together a batch correction was performed. This correction was achieved by calculating the median of all control values (global median). The difference between the global median and the experimental control median was calculated and each measurement of the experiment was increased or reduced by this difference. The results from qPCR for the heatmap and histograms were analyzed using The Mann-Whitney U test.

### Effect Score

The score to estimate relevant effects was calculated by multiplying the –log_10_(p-value) of the Mann-Whitney U test with the hedges effect size and the relative difference of the median. The Mann-Whitney U test was used to estimate, whether the difference between the control and the treatment is significant. The hedges effect size was utilized to obtain a quantitative measure of how strong an effect is considering the spreading of the data points. The relative difference of the median was used to see how different the treatment median is from its respective control.

### Percentage Effect

The Mann-Whitney U test estimates whether the difference between the control and the treatment is significant. Following this, the fold change of the treatment and control is calculated. From this value, percentage effect is calculated. This helps in gaining an estimation of how much treatment varies from the control and there by understand the severity of the effect.

### Literature search

To focus on compounds with poor characterization of possible side effects during embryogenesis in the context of COVID-19 treatment, we performed a systematic literature screen (Fig S8A-B). We retrieved the PubMed ID of each article and counted the total number of articles found. We listed the 15 compounds with most peer-reviewed publications related to COVID-19 and analyzed to which extent these had also already previously been studied in the context of cardiovascular development and embryogenesis (Fig. S8C). We found extensive number of research articles on half of the compounds, and 73.8% of tested drugs had more than ten articles in the cardiovascular system and almost 50% had more than ten articles in embryogenesis (Fig. S8D). However, we found that for approximately half of the compounds studied in the context of COVID-19 there was no literature related to their effect on the cardiovascular system.

The systematic Pubmed search for all investigated compounds was performed using a Jupyter notebook (https://github.com/Alernst/CoVasc_DrugScreen/blob/main/CoVasc_Data_Update.ipynb). The PubchemPy package was used to search synonyms for each compound. These synonyms were combined with specific keywords by “AND” or “OR” operators. Example search term: “Covid-19” AND “Remdesivir” OR “Covid-19” AND “Veklury”. These search terms were generated for each compound combined with context keywords. These search terms were submitted to search in Pubmed using the EntrezPy package. The results were collected as counts and Pubmed IDs. The Pubmed IDs of different context searches were compared for overlapping articles in different contexts.

### Online App

The app was generated in Python code. The code is accessible as a GitHub repository (https://github.com/Alernst/CoVasc_DataApp). The app is hosted via the Streamlit Cloud.

## Acknowledgements

We thank the Medicine For Malaria Venture (MMV) for sharing the *Covid Box*, the animal care takers from EPFL and University of Bern for zebrafish husbandry. The Microscopy Unit from the University of Bern (MIC-Bern) for access to Microscopes and Jochen Gehrig from Acquifer for advise on the Imaging Platform. We are grateful for discussion and advice on the study from Yvonne Döring, Britta Engelhardt, Robert Rieben and the rest of the CoVasc consortium, Beatriz Novoa and Antonio Figueras (IIM-CSIC), Andrew Hemphill (VetSuisse) and comments on the manuscript from Benedetta Coppe.

## Data availability

Results can be found in the data app https://share.streamlit.io/alernst/covasc_dataapp/main/CoVasc_DataApp.py and GitHub https://github.com/Alernst/CoVasc_DrugScreen. All experimental results and scores are in the repository of the data app on GitHub https://github.com/Alernst/CoVasc_DataApp/tree/main/Data.

## Author contributions statement

A.E.: Conceptualization, Methodology, Software, Validation, Formal analysis, Investigation, Data curation, Writing, Visualization.

I.P.: Conceptualization, Investigation, Data curation, Methodology, Formal analysis, Writing, Visualization.

A.M.M.P.: Investigation, Data curation, Methodology, Formal analysis, Visualization, Writing,

N.D.L.: Methodology, Validation, Investigation, Writing.

D.G.: Conceptualization, Methodology, Resources, Supervision, Project administration, Writing

S.L.L.: Conceptualization, Resources, Project administration, Funding acquisition,

A.O.: Resources, Writing.

N.M.: Conceptualization, Methodology, Data curation, Writing, Supervision, Project administration, Funding acquisition.

**Screening for side effects of COVID-19 drug candidates on cardiovascular development Enrst et al.,**

## Supplementary Figure Legends

**Fig. S1.**
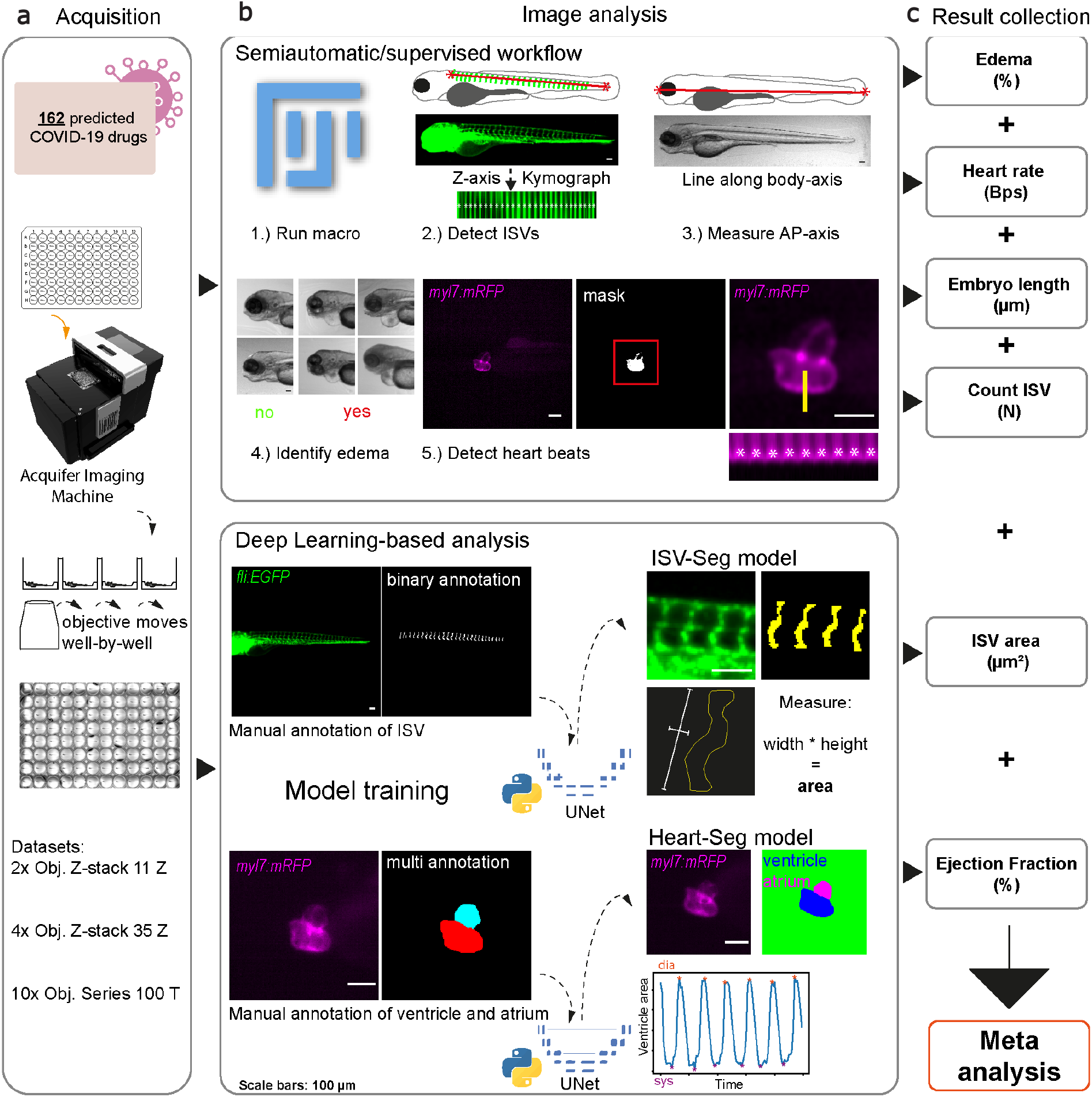
Fluorescent and bright field image segmentation and analysis pipeline. **a** Image acquisition. Treated larvae positioned in a 96 well plate are imaged using the Acquifer Imaging machine. Each well is imaged by a moving objective. **b** Image analysis. (1) In a semiautomatic and supervised workflow based on a Fiji-macro the images are analyzed in a streamlined way but controlled and supported by the user. The user performs the analysis blinded. (2) To count the intersegmental vessels (ISV) the user draws a line (red line) along all ISVs. A kymograph along the Z-axis is generated in which local maxima are detected. The user supervises whether the detection was correct. (3) The larval length from anterior to posterior (AP) is measured by drawing a line from the most anterior of the head to the most posterior of the tail. (4) Effusions are identified as shown in the examples. (5) For heartbeat measurements, hearts are detected with an automated threshold to generate a mask. Subsequently, a kymograph along the time axis is generated to detect the local maxima. The user supervises the detected heartbeats. For the deep learning-based analysis, image data of the ISV and the heart are manually annotated. The annotated data are used to train two different U-Net models. The ISV-Seg model is applied to segment each ISV and calculate the area. The Heart-Seg model is used to segment ventricle and atrium. The minima and maxima of the ventricular area along the time axis are used to calculate the ejection fraction. **c** The results are collected and used for meta-analysis.

**Fig. S2.**
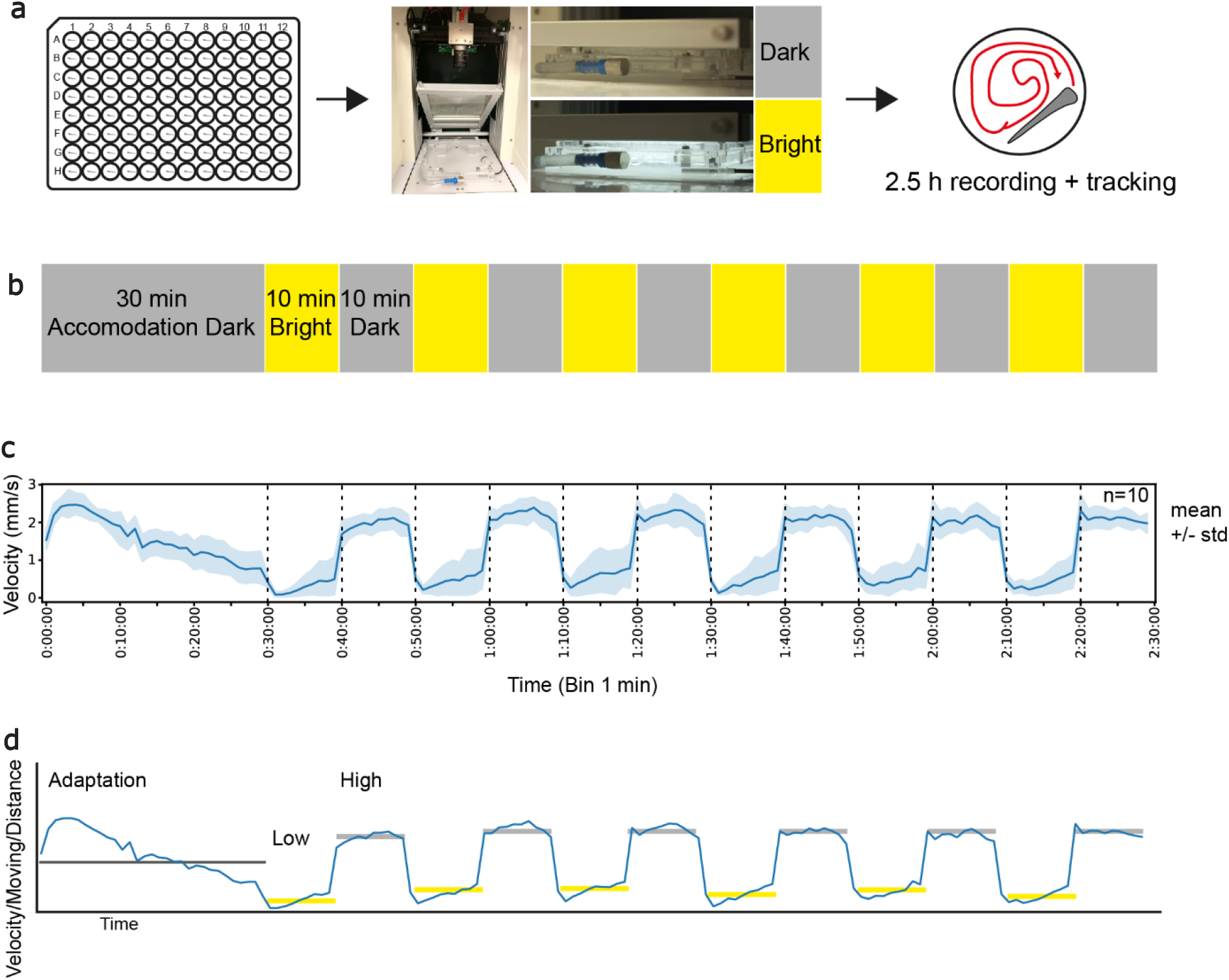
Workflow for behavioral studies in zebrafish larvae. **a** The free-swimming larvae in a 96 well-plate are transferred to the DanioVision recording chamber. The chamber can automatically switch on and off the illumination. Additionally, the larvae are recorded and tracked along 2.5 hours. **b** A bright-dark cycle is applied, starting with 30 min dark for accommodation. Then, the light is switched on for 10 min and switched off again for 10 min. This cycle is repeated in total 6 times. **c** A typical tracking result, here shown as velocity (mm/s) on the X-axis of 10 control larvae over time on the Y-axis (1 min binning, mean +/- standard deviation) is shown with the bright and dark phases (dashed lines). **d** For analysis, the programmed light cycle times are used. The control embryos typically swim less in the bright (yellow line) and more in the dark (grey line).

**Fig. S3.**
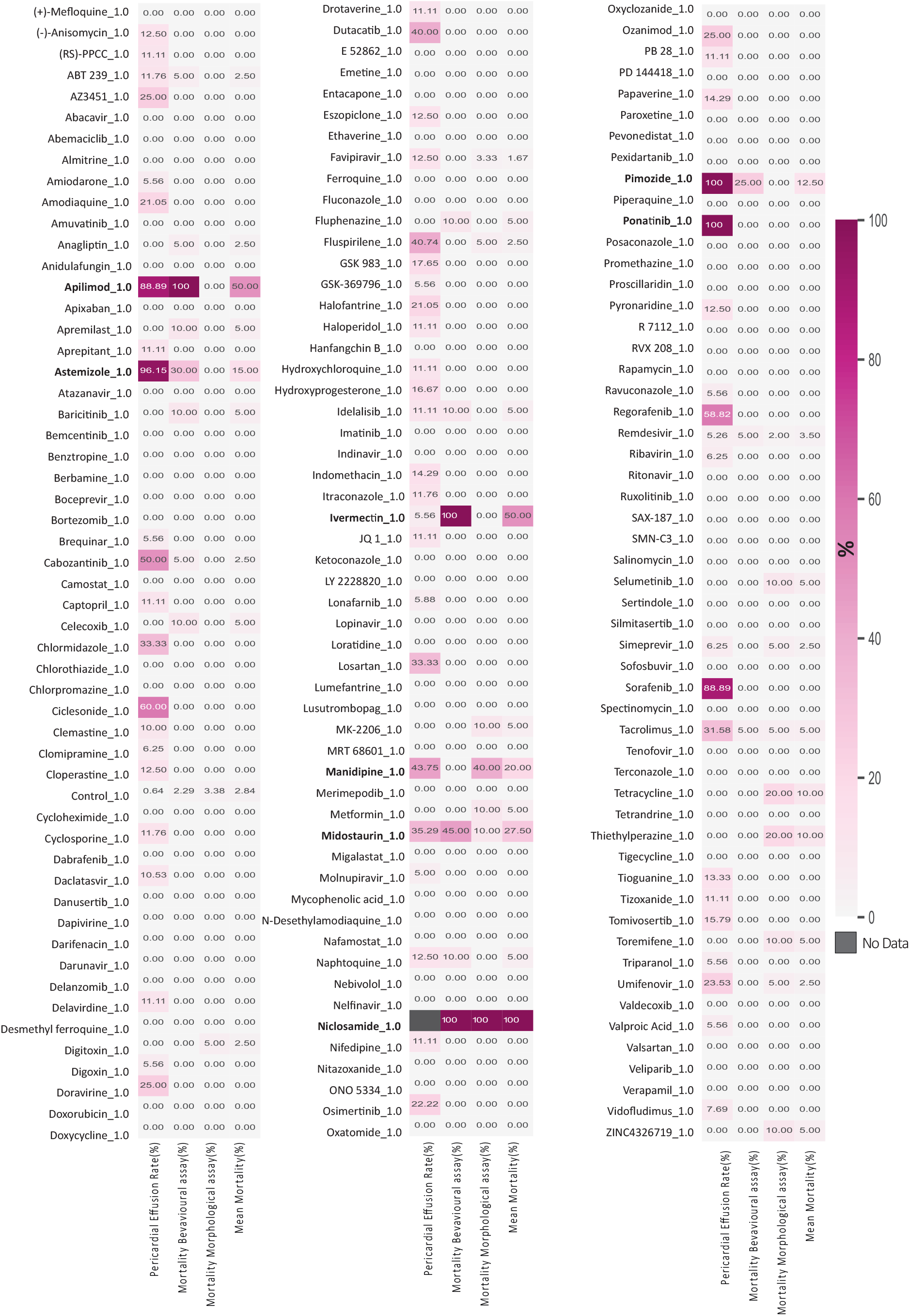
Percentage Mortality and Pericardial Effusion in the drug screen. A heatmap showing Percentage of Pericardial effusion and Mortality in the drug screen. Mortality rates for Morphological Assay(with PTU) and Behavioural Assay (without PTU) are plotted separately followed by a column of mean mortality rate. Dark magenta shows the highest effect.

**Fig. S4.**
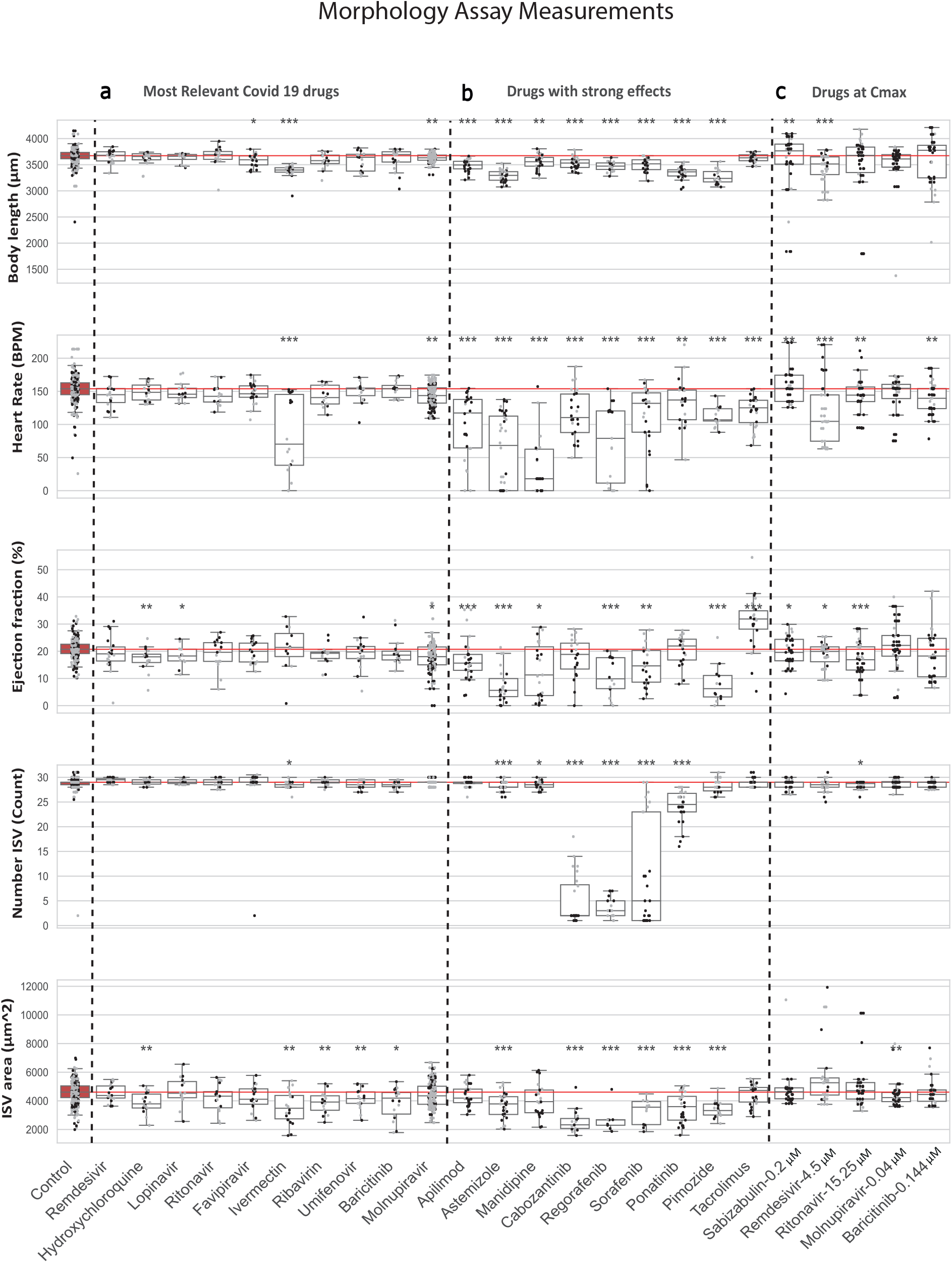
Combined boxplots and swarmplots showing the morphology measurements for each larva. **a** Results from treatments using drugs studied in the context of Covid-19 research. **b** Results from treatments leading to strongest phenotypic alterations. **c** Results from treatments using COVID-19 drugs used at Cmax. The asterisks represent the p-values from the Mann-Whitney U test *<0.05; **<0.01;***<0.001. Results from control larvae highlighted with red fill color. The red line indicates the median of the control as reference for treatment conditions. The plots are shown for all performed measurements

**Fig. S5.**
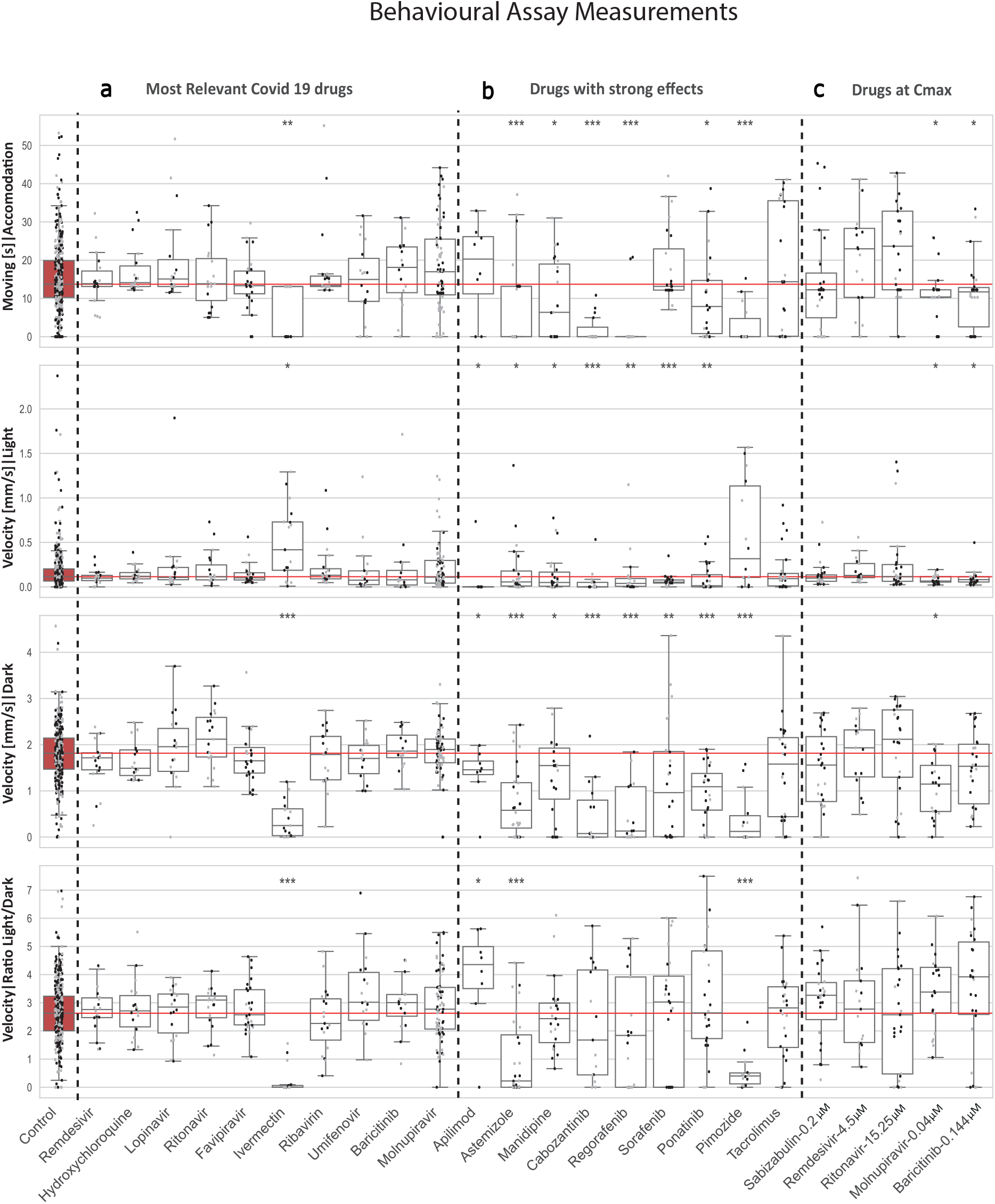
Combined boxplots and swarmplots showing the behavior measurements for each larva. **a** Results from treatments using drugs studied in the context of Covid-19 research. **b** Results from treatments leading to strongest phenotypic alterations. **c** Results from treatments using COVID-19 drugs used at C_max_. The asterisks represent the p-values from the Mann-Whitney U test *<0.05; **<0.01;***<0.001. Results from control larvae are highlighted with red fill color. The red line indicates the median of the control as reference for treatment conditions. The plots are shown for all performed measurements.

**Fig. S6.**
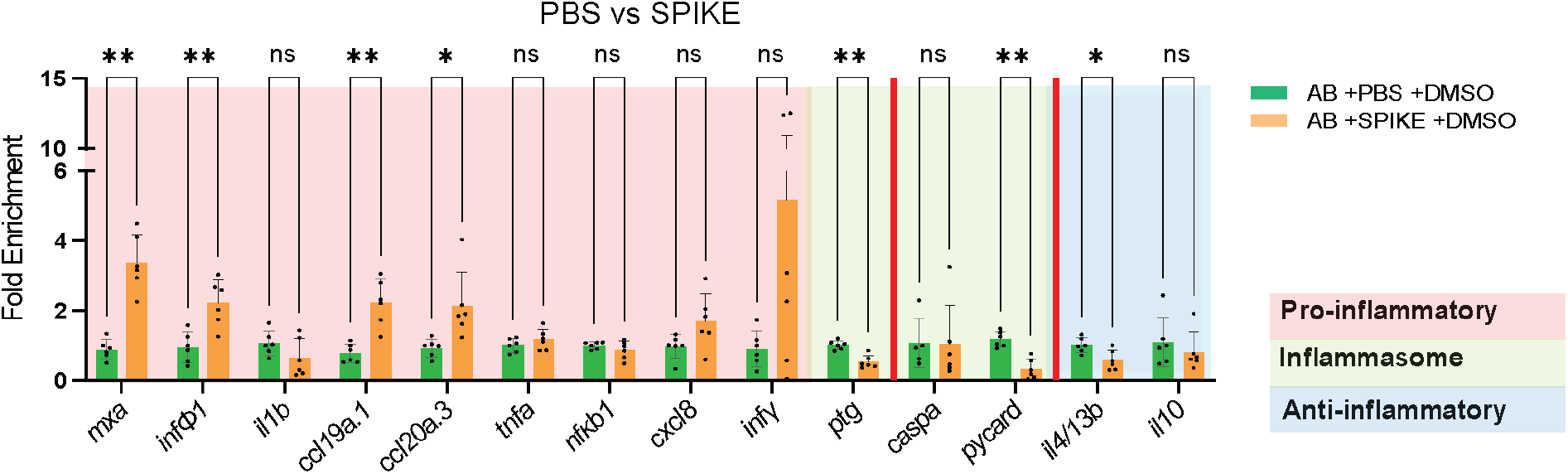
qPCR results for inflammation associated gene expression in zebrafish larvae treated with SPIKE protein in wild type zebrafish larvae. Bars represent mean and SD of 6 biological replicates (pool of 5 embryos each). Asterisks represent p-values from the Mann-Whitney U test *<0.05; **<0.01;***<0.001.

**Fig. S7.**
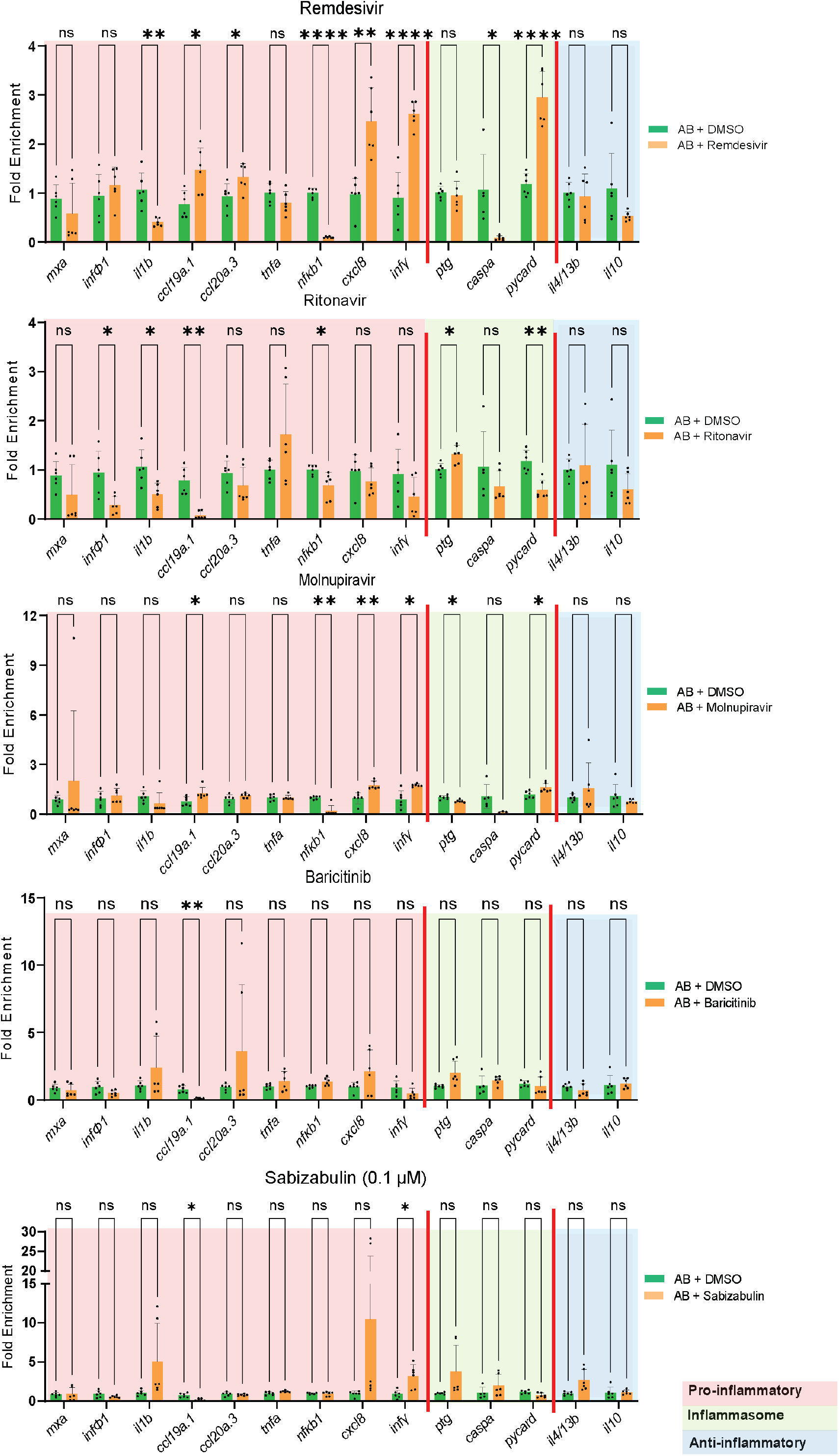
Assessment of immune response upon drug treatment in zebrafish larvae. qPCR results for inflammation associated gene expression in zebrafish larvae treaded with selected compounds. Remdesivir, Ritonavir, Molnupiravir, Baricitinib or Sabizabulin treatment is compared to larvae treated with DMSO. Bars represent mean and SD of 6 biological replicates. Each replicate consists of a pool of 5 larvae. Asterisks represent p-values from the Mann-Whitney U test *<0.05; **<0.01;***<0.001.

**Fig. S8.**
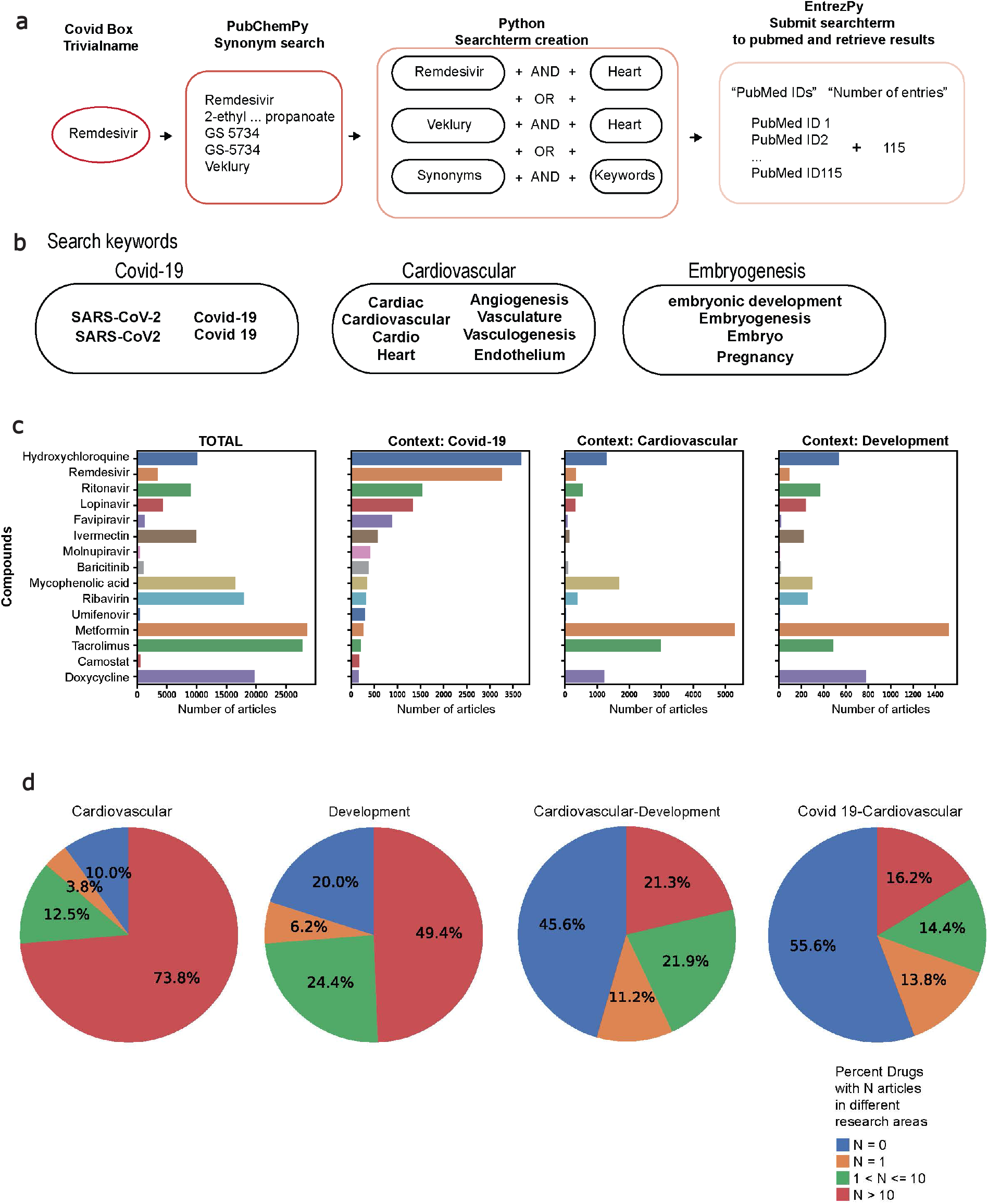
Systematic literature search on published articles describing candidate compounds. **a** Schematic representation of how search term and results were retrieved. **b** In the first bar plot, the 15 compounds with the highest number of publications in the context of COVID-19 are shown. The drug names with black text color were the most mentioned in the literature in the context of COVID-19 plus Molnupiravir and Sabizabulin which was recently approved by several federal agencies for clinical treatment. The additional three plots contain the number of total articles, as well as articles on cardiovascular and embryogenesis research. **c** Pie charts showing the percentage of the compounds (n=162 drugs) with a defined threshold number of articles in the three contexts (Cardiovascular, Embryogenesis, Cardiovascular-Embryogenesis). The search was performed on the 20^th^ of January 2023.

## References

1 Townsend, J. P., Hassler, H. B., Sah, P., Galvani, A. P. & Dornburg, A. The durability of natural infection and vaccine-induced immunity against future infection by SARS-CoV-2. Proc Natl Acad Sci U S A 119, e2204336119, doi:10.1073/pnas.2204336119 (2022).

2 Reardon, S. How well can Omicron evade immunity from COVID vaccines? Nature, doi:10.1038/d41586-022-00283-4 (2022).

3 Wang, Q. et al. Alarming antibody evasion properties of rising SARS-CoV-2 BQ and XBB subvariants. Cell 186, 279–286 e278, doi:10.1016/j.cell.2022.12.018 (2023).

4 MacRae, C. A. & Peterson, R. T. Zebrafish as tools for drug discovery. Nature reviews. Drug discovery 14, 721–731, doi:10.1038/nrd4627 (2015).

5 Rosa, J. G. S., Lima, C. & Lopes-Ferreira, M. Zebrafish Larvae Behavior Models as a Tool for Drug Screenings and Pre-Clinical Trials: A Review. International journal of molecular sciences 23, doi:10.3390/ijms23126647 (2022).

6 Patton, E. E., Zon, L. I. & Langenau, D. M. Zebrafish disease models in drug discovery: from preclinical modelling to clinical trials. Nat Rev Drug Discov 20, 611–628, doi:10.1038/s41573-021-00210-8 (2021).

7 Lee, H. C., Lin, C. Y. & Tsai, H. J. Zebrafish, an In Vivo Platform to Screen Drugs and Proteins for Biomedical Use. Pharmaceuticals (Basel) 14, doi:10.3390/ph14060500 (2021).

8 González-Rosa, J. M. Zebrafish Models of Cardiac Disease: From Fortuitous Mutants to Precision Medicine. Circulation research 130, 1803–1826, doi:10.1161/CIRCRESAHA.122.320396 (2022).

9 Cassar, S. et al. Use of Zebrafish in Drug Discovery Toxicology. Chemical research in toxicology 33, 95–118, doi:10.1021/acs.chemrestox.9b00335 (2020).

10 Gore, A. V., Monzo, K., Cha, Y. R., Pan, W. & Weinstein, B. M. Vascular development in the zebrafish. Cold Spring Harb Perspect Med 2, a006684, doi:10.1101/cshperspect.a006684 (2012).

11 Thomas, L. S. V. & Gehrig, J. Multi-template matching: a versatile tool for object-localization in microscopy images. BMC bioinformatics 21, 44–44, doi:10.1186/s12859-020-3363-7 (2020).

12 Westhoff, J. H. et al. Development of an automated imaging pipeline for the analysis of the zebrafish larval kidney. PloS one 8, e82137–e82137, doi:10.1371/journal.pone.0082137 (2013).

13 Vazao, H. et al. High-throughput identification of small molecules that affect human embryonic vascular development. Proc Natl Acad Sci U S A 114, E3022–E3031, doi:10.1073/pnas.1617451114 (2017).

14 Mauro, A. N. et al. Automated in vivo compound screening with zebrafish and the discovery and validation of PD 81,723 as a novel angiogenesis inhibitor. Sci Rep 12, 14537, doi:10.1038/s41598-022-18230-8 (2022).

15 Maciag, M., Wnorowski, A., Mierzejewska, M. & Plazinska, A. Pharmacological assessment of zebrafish-based cardiotoxicity models. Biomed Pharmacother 148, 112695, doi:10.1016/j.biopha.2022.112695 (2022).

16 Han, Y. et al. Vitamin D Stimulates Cardiomyocyte Proliferation and Controls Organ Size and Regeneration in Zebrafish. Dev Cell 48, 853–863 e855, doi:10.1016/j.devcel.2019.01.001 (2019).

17 Basnet, R. M., Zizioli, D., Taweedet, S., Finazzi, D. & Memo, M. Zebrafish Larvae as a Behavioral Model in Neuropharmacology. Biomedicines 7, doi:10.3390/biomedicines7010023 (2019).

18 Rock, S., Rodenburg, F., Schaaf, M. J. M. & Tudorache, C. Detailed Analysis of Zebrafish Larval Behaviour in the Light Dark Challenge Assay Shows That Diel Hatching Time Determines Individual Variation. Front Physiol 13, 827282, doi:10.3389/fphys.2022.827282 (2022).

19 Jayk Bernal, A., et al. Molnupiravir for Oral Treatment of Covid-19 in Nonhospitalized Patients. N Engl J Med 386, 509–520, doi:10.1056/NEJMoa2116044 (2022).

20 Barnette, G. K. et al. Oral Sabizabulin for High-Risk, Hospitalized Adults with Covid-19: Interim Analysis. NEJM Evidence 1, EVIDoa2200145 (2022).

21 Mandelbaum, J. et al. Zebrafish blastomere screen identifies retinoic acid suppression of MYB in adenoid cystic carcinoma. J Exp Med 215, 2673–2685, doi:10.1084/jem.20180939 (2018).

22 Bose, P. et al. The Novel Small Molecule TRVA242 Stabilizes Neuromuscular Junction Defects in Multiple Animal Models of Amyotrophic Lateral Sclerosis. Neurotherapeutics 16, 1149–1166, doi:10.1007/s13311-019-00765-w (2019).

23 North, T. E. et al. Prostaglandin E2 regulates vertebrate haematopoietic stem cell homeostasis. Nature 447, 1007–1011, doi:10.1038/nature05883 (2007).

24 Lawson, N. D. & Weinstein, B. M. In vivo imaging of embryonic vascular development using transgenic zebrafish. Dev Biol 248, 307–318, doi:10.1006/dbio.2002.0711 (2002).

25 Rohr, S., Bit-Avragim, N. & Abdelilah-Seyfried, S. Heart and soul/PRKCi and nagie oko/Mpp5 regulate myocardial coherence and remodeling during cardiac morphogenesis. Development 133, 107–115, doi:10.1242/dev.02182 (2006).

26 Powrie, Y. et al. Zebrafish behavioral response to ivermectin: insights into potential neurological risk. Medicine in Drug Discovery 16, 100141, doi:https://doi.org/10.1016/j.medidd.2022.100141 (2022).

27 Zhu, X. Y. et al. Ponatinib-induced ischemic stroke in larval zebrafish for drug screening. Eur J Pharmacol 889, 173292, doi:10.1016/j.ejphar.2020.173292 (2020).

28 Jakhar, R., Sharma, C., Paul, S. & Kang, S. C. Immunosuppressive potential of astemizole against LPS activated T cell proliferation and cytokine secretion in RAW macrophages, zebrafish larvae and mouse splenocytes by modulating MAPK signaling pathway. Int Immunopharmacol 65, 268–278, doi:10.1016/j.intimp.2018.10.014 (2018).

29 Patten, S. A. et al. Neuroleptics as therapeutic compounds stabilizing neuromuscular transmission in amyotrophic lateral sclerosis. JCI Insight 2, doi:10.1172/jci.insight.97152 (2017).

30 Oliveira, R., Grisolia, C. K., Monteiro, M. S., Soares, A. M. & Domingues, I. Multilevel assessment of ivermectin effects using different zebrafish life stages. Comp Biochem Physiol C Toxicol Pharmacol 187, 50–61, doi:10.1016/j.cbpc.2016.04.004 (2016).

31 Mosser, E. A. et al. Identification of pathways that regulate circadian rhythms using a larval zebrafish small molecule screen. Sci Rep 9, 12405, doi:10.1038/s41598-019-48914-7 (2019).

32 Sampurna, B. P., Audira, G., Juniardi, S., Lai, Y.-H. & Hsiao, C.-D. A Simple ImageJ-Based Method to Measure Cardiac Rhythm in Zebrafish Embryos. Inventions 3, doi:10.3390/inventions3020021 (2018).

33 Wu, J. Q. et al. A systematical comparison of anti-angiogenesis and anti-cancer efficacy of ramucirumab, apatinib, regorafenib and cabozantinib in zebrafish model. Life Sci 247, 117402, doi:10.1016/j.lfs.2020.117402 (2020).

34 Carra, S. et al. Vandetanib versus Cabozantinib in Medullary Thyroid Carcinoma: A Focus on Anti-Angiogenic Effects in Zebrafish Model. Int J Mol Sci 22, doi:10.3390/ijms22063031 (2021).

35 Chimote, G. et al. Comparison of effects of anti-angiogenic agents in the zebrafish efficacy-toxicity model for translational anti-angiogenic drug discovery. Drug Des Devel Ther 8, 1107–1123, doi:10.2147/DDDT.S55621 (2014).

36 Barac, A. et al. Inappropriate use of ivermectin during the COVID-19 pandemic: primum non nocere! Clin Microbiol Infect 28, 908–910, doi:10.1016/j.cmi.2022.03.022 (2022).

37 WHO. Therapeutics and COVID-19: living guideline, 14 July 2022. Geneva: World Health Organization (2022).

38 WHO. Coronavirus disease (COVID-19): Hydroxychloroquine. (2021).

39 Luo, M., Xie, D., Lin, Z., Sun, H. & Liu, Y. Toxicology evaluation of overdose hydroxychloroquine on zebrafish (Danio rerio) embryos. Sci Rep 12, 18259, doi:10.1038/s41598-022-23187-9 (2022).

40 Drozdzal, S. et al. An update on drugs with therapeutic potential for SARS-CoV-2 (COVID-19) treatment. Drug Resist Updat 59, 100794, doi:10.1016/j.drup.2021.100794 (2021).

41 Mahase, E. Covid-19: Pfizer’s paxlovid is 89% effective in patients at risk of serious illness, company reports. BMJ 375, n2713, doi:10.1136/bmj.n2713 (2021).

42 Roberts, J. A., Duncan, A. & Cairns, K. A. Pandora’s box: Paxlovid, prescribing, pharmacists and pandemic. J Pharm Pract Res 52, 1–4, doi:10.1002/jppr.1799 (2022).

43 Rizk, J. G. et al. Expanded Access Programs, compassionate drug use, and Emergency Use Authorizations during the COVID-19 pandemic. Drug discovery today 26, 593–603, doi:10.1016/j.drudis.2020.11.025 (2021).

44 Hu, W. J. et al. Pharmacokinetics and tissue distribution of remdesivir and its metabolites nucleotide monophosphate, nucleotide triphosphate, and nucleoside in mice. Acta Pharmacol Sin 42, 1195–1200, doi:10.1038/s41401-020-00537-9 (2021).

45 Lingscheid, T. et al. Pharmacokinetics of Nirmatrelvir and Ritonavir in COVID-19 Patients with End-Stage Renal Disease on Intermittent Hemodialysis. Antimicrob Agents Chemother 66, e0122922, doi:10.1128/aac.01229-22 (2022).

46 Painter, W. P. et al. Human Safety, Tolerability, and Pharmacokinetics of Molnupiravir, a Novel Broad-Spectrum Oral Antiviral Agent with Activity Against SARS-CoV-2. Antimicrob Agents Chemother 65, doi:10.1128/AAC.02428-20 (2021).

47 Jorgensen, S. C. J., Tse, C. L. Y., Burry, L. & Dresser, L. D. Baricitinib: A Review of Pharmacology, Safety, and Emerging Clinical Experience in COVID-19. Pharmacotherapy 40, 843-856, doi:10.1002/phar.2438 (2020).

48 Novoa, B. & Figueras, A. Zebrafish: model for the study of inflammation and the innate immune response to infectious diseases. Adv Exp Med Biol 946, 253–275, doi:10.1007/978-1-4614-0106-3_15 (2012).

49 Kraus A., C. E., Huertas M, Chunyan Y., Brafute S., Boudinot P., Levraud J.P., Salinas I.. A zebrafish model for COVID-19 recapitulates olfactory and cardiovascular pathophysiologies caused by SARS-CoV-2. bioRxiv, doi:10.1101/2020.11.06.368191 (2020).

50 Tyrkalska, S. D. et al. Differential proinflammatory activities of Spike proteins of SARS-CoV-2 variants of concern. Sci Adv 8, eabo0732, doi:10.1126/sciadv.abo0732 (2022).

51 Laghi, V. et al. Exploring Zebrafish Larvae as a COVID-19 Model: Probable Abortive SARS-CoV-2 Replication in the Swim Bladder. Front Cell Infect Microbiol 12, 790851, doi:10.3389/fcimb.2022.790851 (2022).

52 Bottiglione, F. et al. Zebrafish IL-4-like Cytokines and IL-10 Suppress Inflammation but Only IL-10 Is Essential for Gill Homeostasis. J Immunol 205, 994–1008, doi:10.4049/jimmunol.2000372 (2020).

53 Chourasia, P. et al. Paxlovid (Nirmatrelvir and Ritonavir) Use in Pregnant and Lactating Woman: Current Evidence and Practice Guidelines-A Scoping Review. Vaccines (Basel) 11, doi:10.3390/vaccines11010107 (2023).

54 Jorgensen, S. C., Tabbara, N. & Burry, L. A review of COVID-19 therapeutics in pregnancy and lactation. Obstet Med 15, 225–232, doi:10.1177/1753495X211056211 (2022).

55 FDA. Subject: Important Safety Information Regarding Use of LAGEVRIO™ (molnupiravir) in Pregnancy and Individuals of Childbearing Potential. (2022).

56 FDA. Sabizabulin Treatment of SARS-CoV-2 Infection in Hospitalized Patients With Moderate to Severe COVID-19 Infection Who Are at High Risk for Acute Respiratory Distress Syndrome (ARDS). November 9, 2022 Meeting of the Pulmonary-Allergy Drugs Advisory Committee (2022).

57 EMA. European Medicines Agency. Covid-19 treatments. https://www.ema.europa.eu/en/human-regulatory/overview/public-health-threats/coronavirus-disease-covid-19/treatments-vaccines/covid-19-treatments (2023).

58 Li, J. T. et al. The mechanism and effects of remdesivir-induced developmental toxicity in zebrafish: Blood flow dysfunction and behavioral alterations. J Appl Toxicol 42, 1688–1700, doi:10.1002/jat.4336 (2022).

59 Coffin, A. B. et al. Putative COVID-19 therapies imatinib, lopinavir, ritonavir, and ivermectin cause hair cell damage: A targeted screen in the zebrafish lateral line. Front Cell Neurosci 16, 941031, doi:10.3389/fncel.2022.941031 (2022).

60 Dinday, M. T. & Baraban, S. C. Large-Scale Phenotype-Based Antiepileptic Drug Screening in a Zebrafish Model of Dravet Syndrome. eNeuro 2, doi:10.1523/ENEURO.0068-15.2015 (2015).

61 Steenbergen, P. J. et al. A Multiparametric Assay Platform for Simultaneous In Vivo Assessment of Pronephric Morphology, Renal Function and Heart Rate in Larval Zebrafish. Cells 9, doi:10.3390/cells9051269 (2020).

62 Anghel, N. et al. Comparative assessment of the effects of bumped kinase inhibitors on early zebrafish embryo development and pregnancy in mice. Int J Antimicrob Agents 56, 106099, doi:10.1016/j.ijantimicag.2020.106099 (2020).

63 Alzahrani, Y., Boufama, B. Biomedical Image Segmentation: A Survey. SN COMPUT. SCI. 2, doi:10.1007/s42979-021-00704-7 (2021).

64 Ronneberger, O., Fischer, P., Brox, T.. U-Net: Convolutional Networks for Biomedical Image Segmentation. Navab, N., Hornegger, J., Wells, W., Frangi, A. (eds) Medical Image Computing and Computer-Assisted Intervention – MICCAI 2015. MICCAI 2015. Lecture Notes in Computer Science 9351, doi:10.1007/978-3-319-24574-4_28 (2015).

65 Kimmel, C. B., Ballard, W. W., Kimmel, S. R., Ullmann, B. & Schilling, T. F. Stages of embryonic development of the zebrafish. Dev Dyn 203, 253–310, doi:10.1002/aja.1002030302 (1995).

66 Lantz-McPeak, S. et al. Developmental toxicity assay using high content screening of zebrafish embryos. J Appl Toxicol 35, 261–272, doi:10.1002/jat.3029 (2015).

67 Vogt, A. et al. Automated image-based phenotypic analysis in zebrafish embryos. Dev Dyn 238, 656–663, doi:10.1002/dvdy.21892 (2009).

68 Zhang, B. et al. Automatic Segmentation and Cardiac Mechanics Analysis of Evolving Zebrafish Using Deep Learning. Front Cardiovasc Med 8, 675291, doi:10.3389/fcvm.2021.675291 (2021).

69 Naderi, A. M. et al. Deep learning-based framework for cardiac function assessment in embryonic zebrafish from heart beating videos. Comput Biol Med 135, 104565, doi:10.1016/j.compbiomed.2021.104565 (2021).

